# Ensemble tests mask missing dynamics in protein conformational generators

**DOI:** 10.64898/2026.09.02.748111

**Authors:** Kaining Liu, Qiuting Qian, Ying Chi

**Affiliations:** Department of Pharmacy of the Second Affiliated Hospital of Zhejiang University School of Medicine, and Zhejiang University–University of Edinburgh Institute (ZJE), Zhejiang University, No. 866 Yu Hang Tang Road, 310058, Zhejiang Province, China; Edinburgh Medical School: Biomedical Sciences, College of Medicine and Veterinary Medicine, The University of Edinburgh, United Kingdom

## Abstract

Protein generators increasingly produce conformational ensembles and trajectories as faster alternatives to molecular dynamics (MD). Yet agreement with an equilibrium ensemble does not reveal how conformations interconvert. We show that ensemble evidence can survive even when temporal dynamics are destroyed. Randomly reordering authentic MD frames leaves ensemble fidelity unchanged but reduces the post-gate kinetic pass rate from 0.963 to 0.004; an independent MD replicate reaches 0.95 under the same test. Dynbench separates conformational coverage, time-dependent behaviour and geometric and fluctuation admissibility. Public generators that appear similar under ensemble tests separate sharply, and in a prospective 36-protein lockbox the reference and floor remain stable while two models reverse order. Ensemble-centred evaluation therefore systematically overstates evidence of learned dynamics. Claims about protein dynamics require direct temporal validation at a resolvable physical interval.

## Introduction

Generative models are moving from producing plausible samples to representing how complex systems evolve. Evaluation has not kept pace: models are often judged by whether their samples reproduce the observed distribution of states. A generator can therefore visit the correct states in the wrong order or at the wrong rates and still appear successful. This mismatch is consequential for proteins. Drug binding, allostery and the opening of hidden pockets depend on both the conformations available and how they interconvert^1^.

Molecular dynamics (MD) simulation supplies a physical clock but remains expensive. Protein generators now sample diverse conformations at far lower cost. Some produce unordered conformations or ensembles^2,3,4,5,6,7,8^; MDGen, ConfRover and newer systems instead produce trajectories whose frame order is intended to carry physical meaning^9,10,11,12,13^. Yet both are commonly judged by equilibrium ensemble agreement. Existing benchmarks assess conformational diversity and plausibility^14^, and experiment-anchored studies test cryptic-pocket populations^15^. These establish which states occur and how often, but the same population can arise from different rates and pathways. Smaller-system studies can test learned evolution operators directly^16^, whereas equilibrium generators accelerate state sampling without making the same temporal claim^17^. Recent latent-space work likewise separates temporal propagation from state generation^18^. Protein generators still lack a shared test of whether their ordered outputs reproduce a measurable transition process.

A simple control exposes this gap. Randomly reordering authentic MD frames preserves every conformation and its frequency while erasing the observed transition sequence. Ensemble tests must return the same answer; a test of dynamics must not. Across 82 proteins, ensemble fidelity is unchanged while time-sensitive tests and the fraction meeting both temporal and geometric criteria collapse almost to zero (Fig. 1). Evidence from the conformational ensemble therefore survives after the detectable dynamics have been destroyed.

**Fig. 1.**
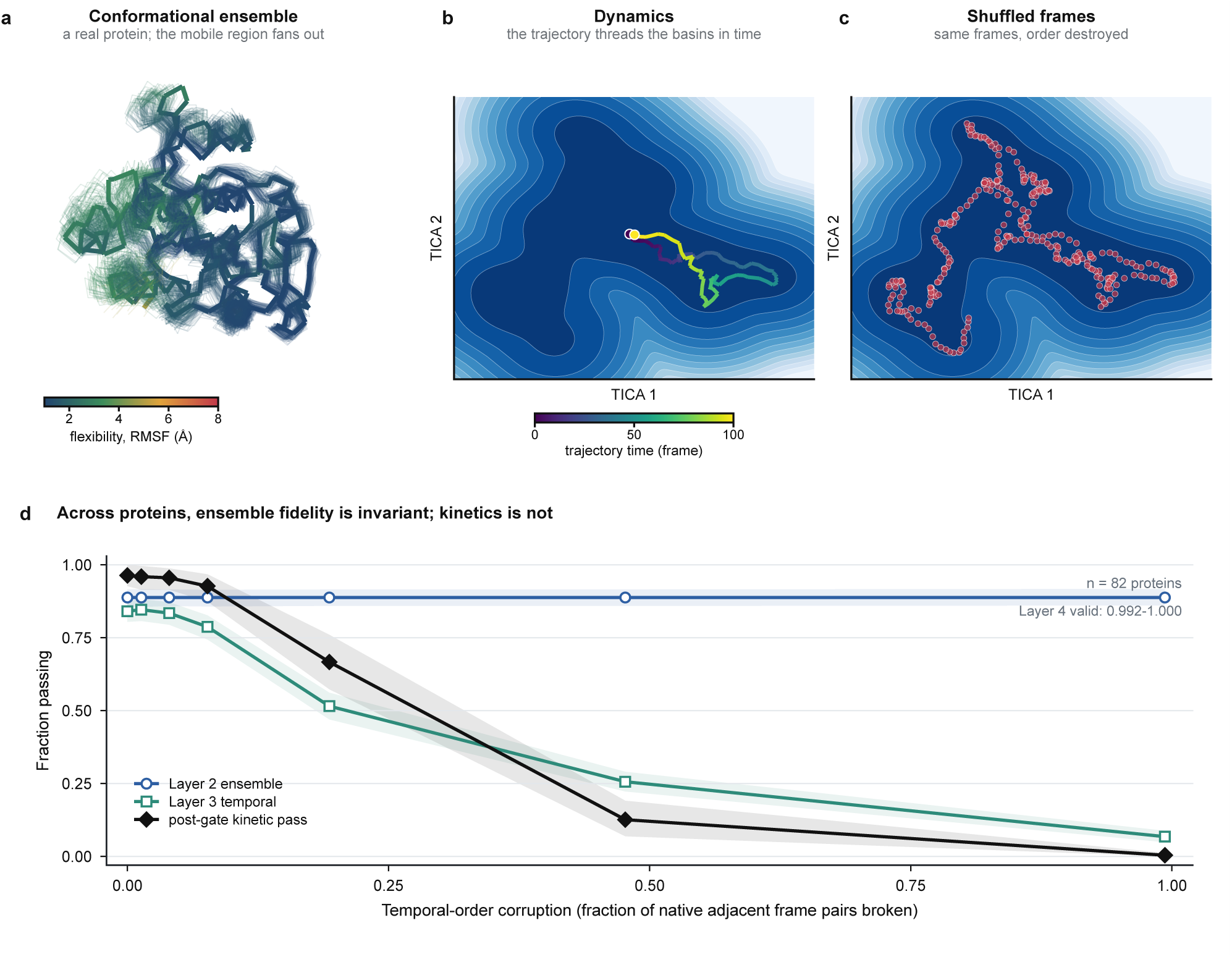
Ensemble fidelity is not dynamical fidelity. One real protein illustrates the distinction, and the frozen cohort establishes that it is not protein-specific. (a) The conformational ensemble of ATLAS 5ftw_A (256 residues), shown as a Cα backbone coloured by per-residue flexibility (root-mean-square fluctuation), with the mobile region fanning into a cloud of overlaid frames. (b) The same trajectory on its time-lagged independent component analysis free-energy landscape, where dark is populated and low in free energy, threading the basins in time order. This is the dynamics. (c) The identical frames with temporal order shuffled. The density is unchanged, yet the ordered process is gone; panels b and c share one landscape and one set of conformations. (d) Controlled temporal-order corruption across all 82 frozen proteins. Every condition contains exactly the same frames. Layer-2 ensemble fidelity is invariant at 0.888 (maximum within-protein change, 0), whereas temporal Layer 3 falls from 0.841 (95% CI, 0.804–0.876) in the intact trajectory to 0.068 (0.049–0.088) after full permutation, and the post-gate kinetic pass rate falls from 0.963 (0.927–1.000) to 0.004 (0.000–0.012). Layer-4 geometric and fluctuation admissibility remains 0.992–1.000. Layer 3 pools informative protein-metric tests, whereas the post-gate rate is the fraction of proteins meeting both the temporal-majority and Layer-4 criteria; the two curves therefore summarize different aggregation levels. Lines show means and shading 95% protein-bootstrap intervals (2,000 resamples) after averaging three fixed permutation seeds within protein; the intact condition is deterministic. Panel d is a temporal ablation; its intact value 0.963 is specific to this ablation setup, not the Table 1 oracle row (0.951).

**Table 1.** Dynbench scorecard on ATLAS-82 (95% bootstrap CIs).

| Model | n | L2 ensemble | L3 temporal | L4 valid | Post-gate kinetic pass |
| --- | --- | --- | --- | --- | --- |
| **Reference controls and comparators** |  |  |  |  |  |
| Held-out MD replicate | 82 | 0.88 (0.85–0.91) | 0.82 (0.78–0.86) | 1.00 (1.00–1.00) | **0.95** (0.89–0.99) |
| Near-harmonic temporal surrogate | 82 | 0.85 (0.81–0.88) | 0.85 (0.81–0.88) | 0.38 (0.28–0.49) | **0.38** (0.28–0.49) |
| Markov-state generator | 82 | 0.81 (0.76–0.86) | 0.29 (0.23–0.36) | 1.00 (1.00–1.00) | **0.32** (0.22–0.43) |
| Independent equilibrium samples | 82 | 0.99 (0.98–1.00) | 0.00 (0.00–0.00) | 1.00 (1.00–1.00) | **0.00** (0.00–0.00) |
| **Ordered models with measurable temporal contracts** |  |  |  |  |  |
| ConfRover | 73 | 0.49 (0.45–0.54) | 0.30 (0.26–0.35) | 0.78 (0.67–0.88) | **0.25** (0.15–0.36) |
| MDGen | 82 | 0.21 (0.17–0.24) | 0.64 (0.60–0.69) | 0.15 (0.07–0.22) | **0.13** (0.06–0.21) |
| **Ordered models with insufficient temporal support** |  |  |  |  |  |
| BioKinema<sup>†</sup> | 81 | 0.64 (0.60–0.68) | 0.13 (0.08–0.18)<sup>‡</sup> | 1.00 (1.00–1.00) | **NR** |
| ProTDyn | 74 | 0.15 (0.12–0.18) | 0.24 (0.17–0.31)<sup>‡</sup> | 0.04 (0.00–0.10) | **NR** |
| MarS-FM | 82 | 0.09 (0.07–0.11) | 0.29 (0.22–0.37)<sup>‡</sup> | 0.00 (0.00–0.00) | **NR** |
| **Unordered ensemble models** |  |  |  |  |  |
| AlphaFlow<sup>§</sup> | 82 | 0.64 (0.57–0.69) | NA | NR | NA |
| P2DFlow | 65 | 0.68 (0.61–0.75) | NA | 0.92 (0.85–0.99) | NA |
| BioEmu | 82 | 0.48 (0.42–0.54) | NA | 0.71 (0.61–0.82) | NA |
| Str2Str | 82 | 0.22 (0.19–0.25) | NA | 0.16 (0.09–0.24) | NA |

L2 and L3 pool informative protein-metric tests meeting their calibrated criterion; L4 and post-gate kinetic pass are per-protein fractions. Parentheses are 95% bootstrap CIs (Methods). *n*, protein coverage. NA means that the output does not define the required task. NR means that the stated temporal interval does not retain enough reference-to-floor separation or informative temporal tests for a stable kinetic estimate. <sup>†</sup>BioKinema uses intended-MSA mode and has 29 of 82 evaluation chains overlapping its training corpus under the model-specific screen (Supplementary Note 3). <sup>‡</sup>Native-interval diagnostic, not used for temporal ranking. <sup>§</sup>AlphaFlow Layer 4 was not evaluated because the complete coordinate batch was unavailable.
The L3 column contains only time-ordered metrics supported at the model's stated interval. Unsupported timescales are not replaced by invented frame-index lags. Detailed calibration, matched-subset analyses and quality-control records are given in Methods and the Supplementary Information.

We use this control as the basis of Dynbench. Each temporal test is compared with an order-destroyed baseline and an independent MD replicate, and the complete generated trajectory is checked for abnormal geometry and fluctuations. The analysis distinguishes limited temporal recovery, inadmissible trajectories and output intervals at which the reference no longer resolves dynamics. A prospective lockbox preserves the distinction between independent MD and the no-dynamics baseline but reverses model order. Together, these results establish time-resolved validation as a separate requirement for claims of learned protein dynamics.

## Results

### Ensemble scores remain high after dynamics are destroyed

Destroying the order of authentic MD frames leaves their ensemble score unchanged. As progressively larger blocks are permuted, ensemble fidelity remains fixed at 0.888 because the conformations and their frequencies are identical. The temporal Layer-3 score falls from 0.841 (95% CI, 0.804–0.876) to 0.068 (0.049–0.088), and the post-gate kinetic pass rate falls from 0.963 (0.927–1.000) to 0.004 (0.000–0.012). Geometric and fluctuation admissibility does not change. Ensemble evaluation is therefore blind to the removal of a temporal process that time-sensitive tests readily detect.

A second control makes the contradiction sharper. Frames drawn independently from the reference ensemble pass 99% of informative ensemble tests, exceeding the ensemble score of a held-out MD trajectory, yet their post-gate kinetic pass rate is 0.00. The construction samples the right conformational distribution but contains no transition sequence. High ensemble fidelity can therefore provide strong evidence for state coverage while providing no evidence for dynamics.

The same separation appears in learned conformational ensembles. We assigned ten arbitrary orders to fixed outputs from BioEmu, P2DFlow and Str2Str^19^, whose official status remained unordered. Their ensemble scores were exactly invariant, whereas their temporal scores remained low across every ordering (0.065–0.158; Extended Data Fig. 5a). A learned state distribution can therefore retain its apparent quality under many incompatible temporal stories.

These controls expose the same blind spot at increasing levels of realism. Frame permutation preserves structures and populations while removing their observed sequence. Independent sampling removes all serial dependence, and arbitrary orderings show that one learned ensemble can support many incompatible trajectories. Ensemble tests locate generated structures and estimate their populations; they do not identify the process that moves probability between them.

### Independent MD establishes a measurable temporal target

Independent simulations establish that the temporal signal is measurable. A held-out MD replicate, evaluated as an ordinary trajectory, reaches a post-gate kinetic pass rate of 0.95 (0.89–0.99), while the independently sampled floor remains at 0.00. Under the same 1-ns measurement, the reference data contain a reproducible temporal signal that disappears when frame order is destroyed.

Independent MD provides a positive reference, whereas independent equilibrium samples provide a no-dynamics baseline. The replicate measures temporal agreement that survives stochastic variation between simulations; the baseline retains the population without a transition process. Their separation shows that the target is measurable and that equilibrium agreement is insufficient to reach it. The fraction of this difference recovered by a model provides an interpretable measure of its temporal evidence.

This separation is stable across reference replicates. Rotating the held-out replicate keeps the post-gate kinetic pass at 0.951–0.976 while the independently sampled baseline remains at 0.00; only 1.9% of metric-by-protein cells lack a resolvable target. Nearly all tested measurements therefore distinguish independent MD from samples in which temporal order has been removed.

Exact-frame shuffling asks whether temporal order matters at all. A harder control asks whether temporal statistics respond when every conformation remains fixed but the transition process changes. Across the same 82 proteins, a reversible state model reordered an identical coordinate multiset under five nominal dynamical-speed factors. In the two pre-specified extreme contrasts, the 0.25-fold arm increased temporal Layer 3 by 0.067 (95% CI, 0.046–0.090), whereas the 4-fold arm reduced it by 0.136 (0.107–0.164). Post-gate pass moved in parallel, by +0.085 and −0.183. Layer 2 remained fixed at 0.990 and Layer 4 within 0.003. Dynbench therefore detects graded transition-rate perturbations while ensemble and geometric quality remain fixed (Extended Data Fig. 7).

This observation determines the design of Dynbench (Fig. 2). The benchmark first asks whether an output reproduces the reference conformational ensemble. For ordered trajectories, it then tests autocorrelation, implied timescales, directional predictability and state-transition statistics only where independent MD distinguishes them from the order-destroyed baseline. Finally, it checks the complete trajectory for abnormal geometry and fluctuations. A post-gate kinetic pass requires both temporal evidence and an admissible trajectory. Unordered ensembles retain their ensemble result but receive no temporal verdict because no time axis was claimed.

**Fig. 2.**
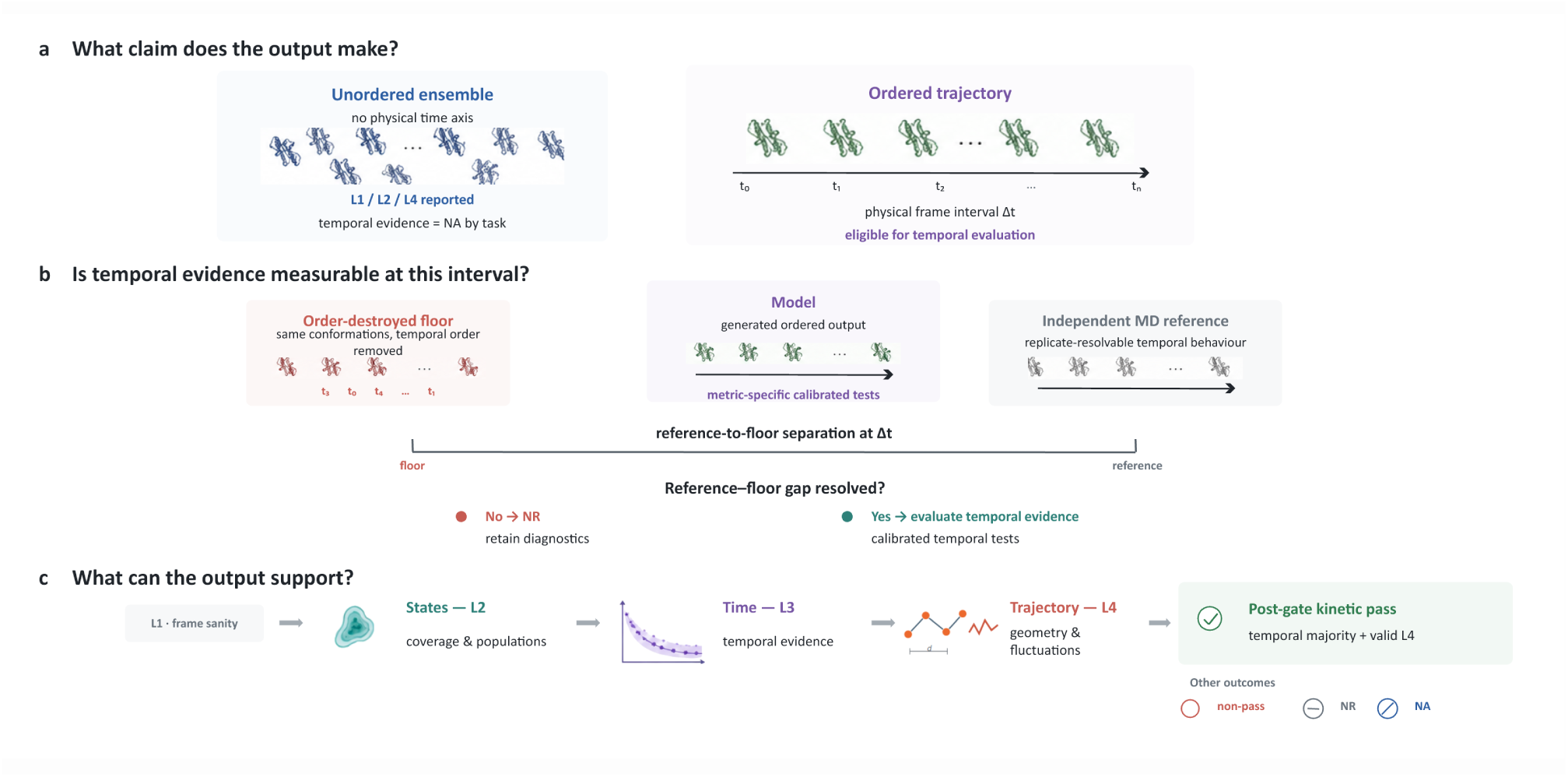
Dynbench separates output claims, temporal measurability and the evidence they can support. Schematic; no data. (a) Unordered ensembles report frame, ensemble and trajectory-quality evidence but make no temporal claim, whereas ordered trajectories declare a physical frame interval and are eligible for temporal evaluation. (b) At that interval, temporal evidence is interpreted only when an independent MD reference can be distinguished from an order-destroyed floor. An unresolved reference-to-floor gap yields NR rather than a model failure. (c) Resolved outputs are interpreted separately through frame sanity, state agreement, calibrated temporal evidence and trajectory-level geometric and fluctuation admissibility. A post-gate kinetic pass requires both a majority of informative temporal tests and valid Layer 4; resolved non-pass, NR and NA remain distinct outcomes. Detailed metrics, verdict definitions and thresholds are given in Methods.

The reference controls reveal behaviours that a single score would merge. A near-harmonic temporal surrogate reaches a post-gate kinetic pass rate of 0.38 (0.28–0.49) by reproducing several relaxation statistics, but often leaves the region of admissible protein motion. A Markov-state generator reaches 0.32 (0.22–0.43) by recovering transitions between coarse states while losing details of the conformational distribution. Similar final scores can therefore encode different scientific outcomes. Separating state coverage, temporal statistics and geometric and fluctuation admissibility preserves those distinctions.

The central contrast persists under a different temporal reference. When every temporal metric is judged against the held-out MD replicate, the reference and floor remain at 1.00 and 0.00. ConfRover moves from 0.25 to 0.40, whereas MDGen remains near 0.13. The measurement gap is stable, while the absolute position of an intermediate model depends on the chosen scientific ruler.

### Public generators differ in temporal signal and trajectory quality

P2DFlow, BioEmu, AlphaFlow and Str2Str generate conformations without assigning them a physical time axis, so their kinetic result is NA. P2DFlow, for example, reaches ensemble fidelity 0.68 (0.61–0.75) and geometric and fluctuation admissibility 0.92 (0.85–0.99), supporting its role as an ensemble generator while leaving kinetics unassessed. ATMOS was not independently scored because public code and weights were unavailable at the evaluation cutoff; its reported 1-ns results are included in the literature comparison.

Second, the two ordered models with measurable 1-ns outputs show different limitations. ConfRover passes 0.30 of informative temporal tests and remains geometrically admissible on 0.78 of proteins, producing a post-gate kinetic pass rate of 0.25. MDGen passes a larger temporal fraction of 0.64, but remains admissible on only 0.15 of proteins and falls to a post-gate kinetic pass rate of 0.13. A promising temporal statistic can therefore occur in a geometrically inadmissible trajectory. Neither model approaches the held-out MD reference of 0.95.

Temporal or structural evaluation alone would obscure this contrast. ConfRover retains a smaller temporal signal but produces admissible trajectories more consistently. MDGen more often reproduces the tested temporal statistics, yet those statistics frequently occur in trajectories that fail the geometric or fluctuation guardrails. The two models therefore require different improvements: stronger temporal propagation for ConfRover and more coherent trajectories for MDGen.

A same-protein, reference-assisted diagnostic localises this loss. Under a common 100-frame comparison, the independent-MD reference and no-dynamics baseline define the available range. Rescaling fluctuations and correcting virtual bond lengths raises the share of this range recovered by 9 percentage points for ConfRover and 24 for MDGen. Four of five ConfRover rescues and 17 of 19 MDGen rescues arise solely from crossing the geometric gate rather than from improved temporal statistics (Extended Data Fig. 3). Most rescued proteins already carried temporal evidence, but their complete trajectories were inadmissible. MDGen therefore contains more recoverable temporal structure than its headline pass rate suggests, delivered with poorly calibrated fluctuations or chain geometry.

Applying the same calibration to the model, independent-MD reference and no-dynamics baseline avoids crediting a shift in the reference to the model. On 73 shared proteins, the increase is larger for MDGen than for ConfRover by 0.181 (95% CI, 0.062–0.303). Because it uses same-protein reference MD, this analysis localises geometry-related losses.

### Some output clocks cannot support a kinetic claim

Third, three ordered outputs remain kinetically unresolved at their stated intervals. BioKinema^20^ and ProTDyn retain too little reference-to-floor range at 10 ns, while the approximately 50-ns MarS-FM reference contains only three frames and a median of one informative temporal test. Their native-interval values are reported as diagnostics rather than ranked kinetic results. This classification is unchanged across all 15 tested combinations of range and support thresholds. A reference-only analysis further shows that temporal resolution and observation horizon act jointly: increasing the interval without extending the trajectory eventually removes the evidence needed to distinguish intact from order-destroyed dynamics (Extended Data Fig. 5b).

NR identifies reference sampling that cannot support a kinetic estimate because intact and independently sampled references have become nearly indistinguishable.

Geometric results remain informative even when temporal interpretation is unavailable. BioKinema passes the geometric and fluctuation guardrails on all 81 covered proteins, whereas ProTDyn passes on 3 of 74 and MarS-FM on none. MarS-FM expands globally on 74 of 82 proteins, with a median radius-of-gyration ratio of 5.42. These results distinguish a geometrically stable output with an unresolved time interval from a trajectory that fails the geometric criteria outright (Extended Data Figs. 1 and 2).

### Prospective data preserve the measurement but reverse model rank

A prospective 36-protein lockbox tests whether the measurement survives outside the dataset on which the main scorecard was developed (Fig. 4). With three independent 200-ns reference simulations, held-out MD and the order-destroyed floor remain cleanly separated at 1.00 and 0.00. The models change order. MDGen reaches a post-gate kinetic pass rate of 0.58 and ConfRover 0.11, a paired difference of −0.47 (95% CI, −0.67 to −0.25). MDGen leads on both the temporal and geometric components, locating the reversal in both parts of the scorecard. The direction and reference separation persist after removing nine proteins flagged against disclosed training corpora and after removing each temporal metric family in turn.

**Fig. 3.**
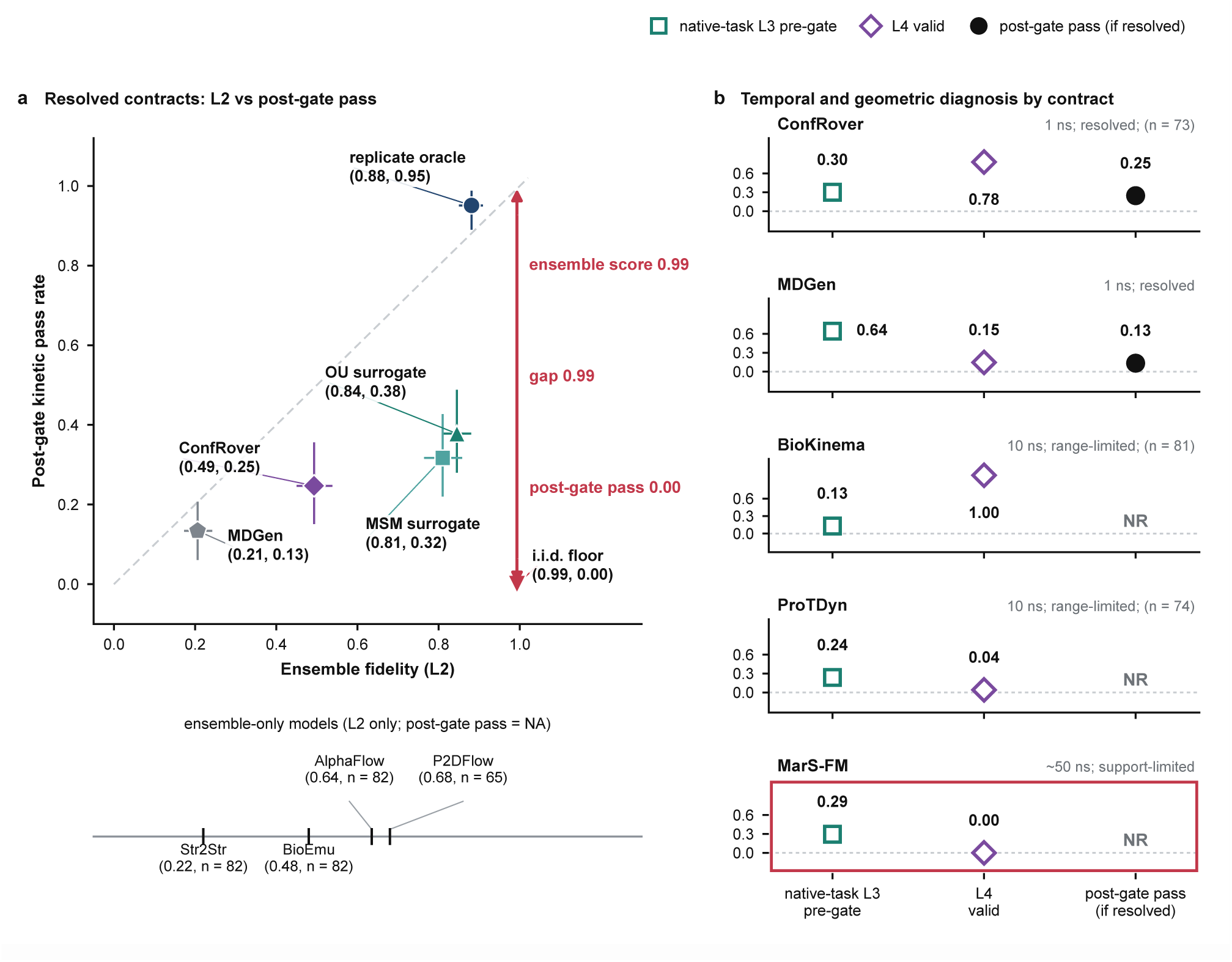
Ensemble agreement and temporal evidence lead to different conclusions. (a) Ensemble fidelity versus post-gate kinetic pass for the fixed 1-ns reference controls, comparators and ordered models whose temporal contracts are measurable. Error bars are 95% protein-bootstrap CIs. Independent equilibrium samples occupy the high-ensemble, zero-kinetics corner, whereas held-out MD passes the post-gate test. Unordered models are shown on a separate ensemble rail because they make no temporal claim. The horizontal axis pools informative protein-metric tests, whereas the vertical axis is a per-protein binary fraction; distances in this plane are descriptive and do not define a composite performance metric. (b) Temporal and geometric results for five ordered models at their stated output intervals. A post-gate kinetic pass is shown only when the matching reference retains sufficient temporal information; the remaining native-interval values are retained as diagnostics.

**Fig. 4.**
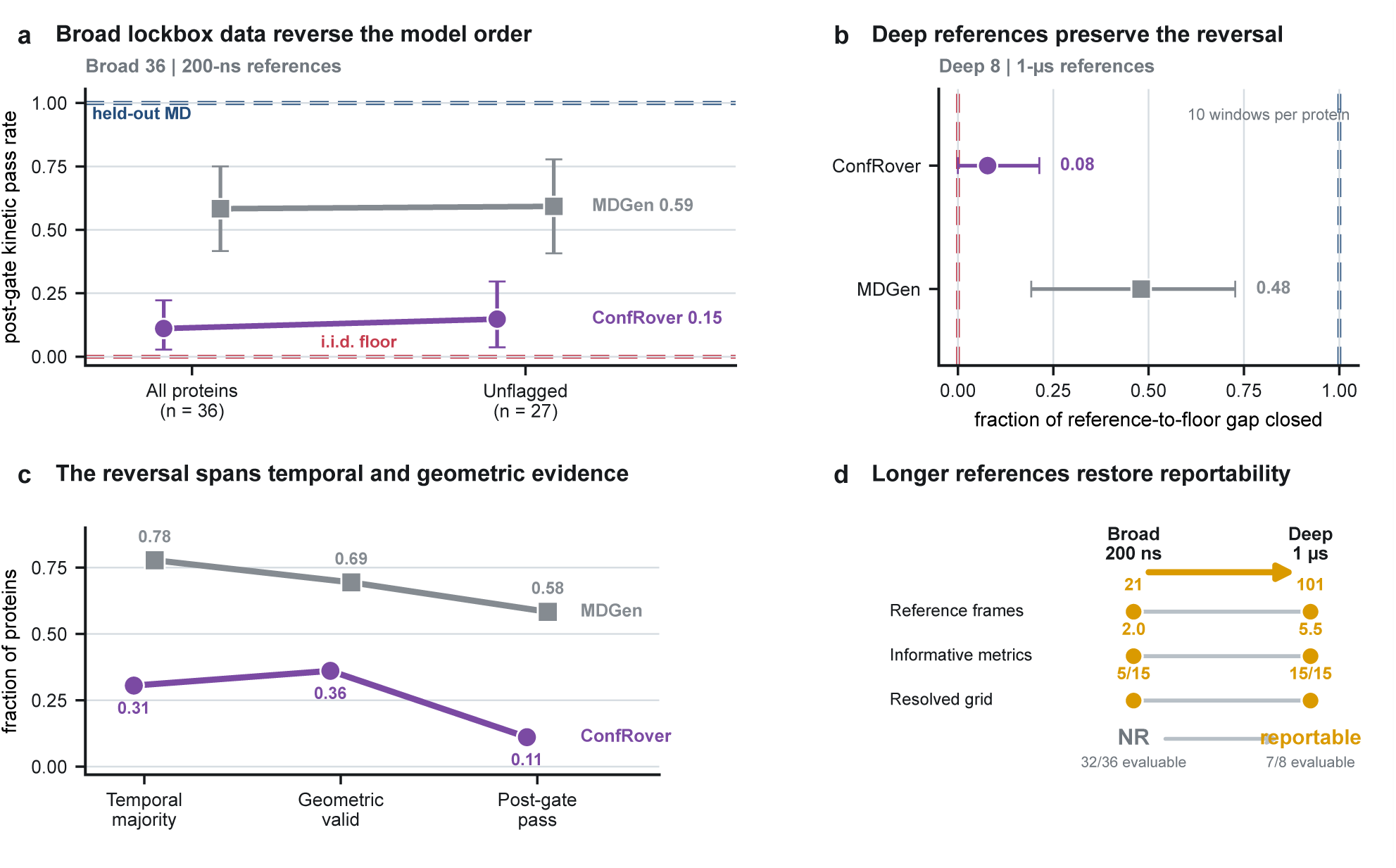
Prospective Lockbox-36 validation preserves the measurement while changing model order. (a) Post-gate kinetic pass rates for ConfRover and MDGen on all 36 proteins and after removing nine proteins flagged by the post-result sequence-overlap screen. Dashed lines mark the held-out-reference and independently sampled anchors; error bars are 95% protein-bootstrap intervals from 2,000 draws. (b) Fraction of the reference-to-floor gap closed on eight preselected proteins after extending all three reference replicates to 1 μs. (c) Temporal, geometric and combined results show that MDGen’s lead is present in both components of the scorecard. (d) Extending the BioKinema reference from 200 ns to 1 μs increases the number of reference frames and informative temporal tests, making its 10-ns output measurable. Missing model outputs are treated as technical NA rather than failures. Source data are provided with this paper.

BioEmu remains an ensemble model, reaching ensemble fidelity 0.813 with no temporal result.

The independent-MD reference and order-destroyed baseline remain stable even though the models change order. The measurement therefore transfers more reliably than the ranking. ConfRover’s advantage is specific to ATLAS, whereas MDGen gains across both temporal and geometric measurements on the lockbox. Relative model performance thus depends on the protein distribution even when the temporal signal remains measurable.

Longer reference simulations on eight preselected proteins provide a second prospective test. After extending each replicate to 1 μs, MDGen recovers 48% of the difference between independent MD and the no-dynamics baseline, compared with 8% for ConfRover. The longer references also make BioKinema’s 10-ns output measurable. Seven of eight proteins are technically evaluable, temporal support rises from a median of 2.0 to 5.5 tests and all 15 reportability settings are resolved. Yet BioKinema passes on only 1 of 7 proteins in the primary sample and on none in four additional samples. More reference data can therefore turn an unanswerable question into an answer without turning limited model evidence into success.

### Independent datasets and positive controls reproduce the measurement gap

The ensemble-temporal separation extends beyond ATLAS. On 82 mdCATH domains^21^, held-out MD reaches 1.00, ConfRover reaches 0.07 and the independently sampled floor remains at 0.00. ConfRover exceeds the floor by 0.07 (95% CI, 0.02–0.13) but remains 0.93 below the reference. A separate set of 85 phosphorylated proteins retains the same anchor pattern, with temporal scores of 0.87 for the replicate and 0.00 for the floor. The mdCATH arm is a qualitative external replication under a replicate-based ruler; its absolute score is on a different scale from the ATLAS scorecard.

Fast peptide simulations provide a complementary positive control where kinetics are well sampled^9,22,23^. Held-out trajectories from two alanine-dipeptide sources^22,24^ reach a post-gate kinetic pass rate of 1.00 while their shuffled floors remain at 0.00; the five-residue WLALL system yields the same separation across 25 independent trajectories.

The central contrast persists across temporal thresholds, alternative held-out replicates, leave-one-protein-out analyses and alignment-screened subsets (Extended Data Fig. 4). Removing any protein moves no post-gate row by more than 0.014. Removing the 10 proteins flagged by the alignment screen changes no row materially and shifts the reference-to-floor separation by −0.004 (95% CI, −0.013 to 0.000).

## Discussion

Protein generators can now produce convincing collections of conformations, but a collection of states is not yet a model of motion. Our central experiment makes that distinction empirical. The same authentic MD frames retain exactly the same ensemble score after their temporal order is destroyed, while the kinetic signal collapses. Independent simulations recover the signal under the same measurement. Ensemble agreement can therefore certify the states present in an output without testing the process that connects them.

The gap is a problem of identifiability: an equilibrium distribution does not determine the transition operator that generated it, and distinct pathways and rate scales can preserve the same state populations^25^. Unordered generators can therefore be strong ensemble models without making a temporal claim. Ordered generators require a declared physical interval, reference data that resolve that interval and trajectories that remain admissible as they evolve.

Ensemble benchmarks and experiment-anchored population measurements remain essential because they test whether the relevant conformations are sampled and weighted correctly^14,15^. Dynbench addresses the complementary question of whether generated trajectories connect those conformations with measurable temporal behaviour. A model must satisfy both requirements before its output can support a claim about protein dynamics.

The model results make these distinctions concrete. On ATLAS, ConfRover usually maintains admissible trajectories but recovers only a modest fraction of the measured temporal behaviour. MDGen reproduces more of the tested autocorrelation and transition statistics, yet its full trajectories often fail the fluctuation or chain-geometry criteria. Reference-assisted calibration shows that much of this temporal signal is already present and becomes visible when those specific defects are reduced. BioKinema presents a third case: its 10-ns output cannot be judged with the shorter reference, but extending the same proteins to 1 μs makes the test measurable and reveals limited temporal evidence. These outcomes point to different problems in temporal propagation, trajectory quality and reference duration; a single aggregate rank would conceal them.

The prospective lockbox separates reproducibility of the measurement from reproducibility of model rank. Independent MD remains distinct from the no-dynamics baseline while ConfRover and MDGen reverse order. Extending the reference trajectories also turns BioKinema from an unresolved output interval into a measurable test with limited model evidence. These results distinguish whether temporal behaviour is measurable, whether a model contains it and whether the resulting trajectory remains usable.

Dynbench provides a practical test of whether an ordered protein generator reproduces measurable temporal behaviour. Exact-frame shuffling reveals when ensemble evidence contains no information about order, independent MD establishes that the missing signal can be resolved, and the geometric criteria identify favourable temporal statistics carried by failing trajectories. Ensemble generators can be reported through state coverage and population agreement. Ordered generators additionally require a physical frame interval, reference data that retain temporal information and a complete trajectory that remains admissible. Reporting these results separately shows whether a limitation lies in temporal behaviour, trajectory quality or the reference sampling itself.

Dynbench resolves Cα-level geometry and fluctuations, primarily at 1-ns sampling over approximately 100 ns, with longer prospective references on a smaller set. Energetic validity, slower biological rates and broader model ordering require matched experimental or simulation support. Within the measured regime, the separation between intact references and order-destroyed controls is large, reproducible across datasets and preserved under the principal sensitivity analyses.

Protein generation has progressed from predicting single structures to sampling conformational populations. The next step is not merely to generate more plausible states, but to show that those states are connected by the right measurable process. Claims that a generator has learned protein dynamics should therefore require independent temporal evidence at a resolvable physical interval, not ensemble realism alone.

## Methods

### Reference data and split

Reference trajectories are the all-atom MD ensembles of ATLAS^26^. Each protein provides three independent 100-ns replicas saved every 10 ps. The anchor ladder uses 1-ns-spaced frames, so its temporal result concerns 1-ns-interval kinetics within a ∼100-ns window. Deep-model trajectories retain their native or intended interval: 1 ns for ConfRover and MDGen, 10 ns for BioKinema and ProTDyn, and approximately 50 ns for MarS-FM. The temporal-support audit maps every requested lag to physical time and marks unsupported metrics NA. The anchor bracket is therefore a fixed-resolution experiment, whereas deep-model temporal values follow their stated contracts unless a common physical interval is available. The fixed evaluation set contains 82 protein chains. Four models have partial coverage: ConfRover 73, ProTDyn 74, P2DFlow 65 and BioKinema 81. Because the ATLAS trajectories are public, this evaluation is not a strict hidden test. A post hoc MMseqs2^27^ audit at ≥30% identity and ≥50% query coverage tests robustness to sequence overlap in the auditable shared corpus; complete model-specific training-corpus disjointness remains unverified.

Frozen release protocol. Dynbench froze the official split, anchor definitions, per-metric verdict thresholds, bootstrap specification, Layer-4 decision logic and applicability-by-task rules before model promotion. The study uses a version-frozen protocol rather than prospective preregistration; exploratory scoring preceded the split freeze, and later analyses are identified as post hoc.

### External protein datasets

Two further protein datasets test generality off ATLAS. mdCATH^21^ provides multi-temperature all-atom MD for CATH domains, with ∼440-ns replicas saved every 1 ns, compared with 100 ns for ATLAS. We score 82 domains at 320 K under a 4-ns contract and take a held-out temperature-matched replicate as the oracle. The ConfRover submission records a 4-ns frame interval and 110 model frames; the five reference replicates are evaluated at the same interval and contain 110–113 frames, as recorded per protein by the time-alignment audit. This bounded external arm evaluates one ConfRover contract, for which model-specific training-corpus disjointness is unverified. PTM-85 is a set of 85 phosphorylated proteins (phospho-serine) with three independent 10 ns MD replicates each, Cα read from the atom14 slot-1 convention, the oracle a held-out replicate. It is used for the baseline ladder. Both datasets use the same four-layer implementation, with source, stride, observable and oracle/floor definitions recorded for reproducibility.

### Prospective Lockbox-36 validation

Lockbox-36 contains 36 soluble monomeric proteins of 60–300 residues selected prospectively after the evaluated model checkpoints were fixed. Protein identities, three replicate roles, the eight-protein Deep subset, model contracts, endpoints, exclusions, bootstrap rules and gate thresholds were frozen before model scoring. Each Broad protein has three independent 200-ns CHARMM36m simulations in modified TIP3P water with 150 mM NaCl, saved every 10 ps; the three replicas of each preselected Deep protein were extended to 1 μs. Whole molecules were reconstructed across periodic boundaries before Cα extraction, and the primary analysis used a 1-ns interval. The last replicate was the evaluation target, the first two formed the fit and i.i.d.-floor pool, and the second served as the reference stand-in. Floors used 20 seeds. Broad estimates use the first 100 aligned frames; Deep estimates average ten non-overlapping 100-frame windows within each protein before protein-level aggregation. ConfRover and MDGen are the primary ordered 1-ns models, BioEmu is an unordered applicability control and BioKinema is a secondary 10-ns contract audit. Four BioKinema Broad proteins and one Deep protein are technical NA rather than performance failures. Confidence intervals use 2,000 protein bootstrap draws, with windows averaged within protein. A post-result MMseqs2 sensitivity at 30% identity and 50% query coverage flags 9 of 36 proteins against the disclosed ATLAS or mdCATH corpora; removing them does not change the paired model direction. A post hoc metric-family sensitivity removes the timescale, autocorrelation, transition/MFPT or directional/VAMP family as a complete group, recomputes temporal majority and reportability from the stored verdicts, and uses the same paired protein bootstrap with crossed floor-seed resampling. The family definitions were formalised after scoring and no metric combinations were searched. The complete analysis and audit records are included in the public evidence package.

### Auditing the data behind dynamics claims

To test whether the protocol detects replicate-resolvable kinetics and, separately, whether simple models reproduce them, we ran the trivial-floor and replicate-oracle construction on small-peptide MD with no generative model in the loop. Two alanine-dipeptide sources were used: the Timewarp release^22^ (two independent trajectories, 0.25 ps saving) and the MDshare heavy-atom set distributed with PyEMMA^24^ (three independent 250 ns trajectories, 1 ps saving). An intermediate five-residue point is provided by the MDshare WLALL pentapeptide set, comprising 25 independent 500-ns implicit-solvent trajectories loaded at stride 25. This set furnishes the strongest cross-replicate peptide oracle in the study. On the pentapeptide the Layer-3 kinetic axis has a single informative metric (7 of its 8 Layer-3 metrics saturate on this simple system and are excluded; the surviving one is time-ordered, so it is the kinetic gate that it carries). On it the replicate oracle, the MSM generator and the OU surrogate all pass (post-gate kinetic pass 1.00) while the i.i.d. floor fails (0.00). This is the intermediate positive control of Fig. 5, run CPU-only with no generative model. Peptides of two to five residues have too few Cα atoms for the protein featurisation, so their kinetic observable is the full set of heavy-atom pairwise distances. The trivial floor, replicate-calibrated oracle, TICA basis, implied-timescale and autocorrelation statistics and the kinetic verdict are otherwise identical to the protein pipeline. Independent trajectories furnish the oracle directly as a held-out replicate. Trajectories are subsampled at a stated stride chosen to retain the system’s intrinsic timescale. Provenance records specify the public source, stride, observable and oracle/floor definitions. For model rows gated by Layer 4, a separate geometry-evidence record reports per-protein virtual-bond statistics, the invalid-protein fraction and an extraction sanity check. The latter is the population-median Cα–Cα bond, which must sit near 3.8 Å. These records distinguish a genuine geometric-admissibility failure from an adapter error.

**Fig. 5.**
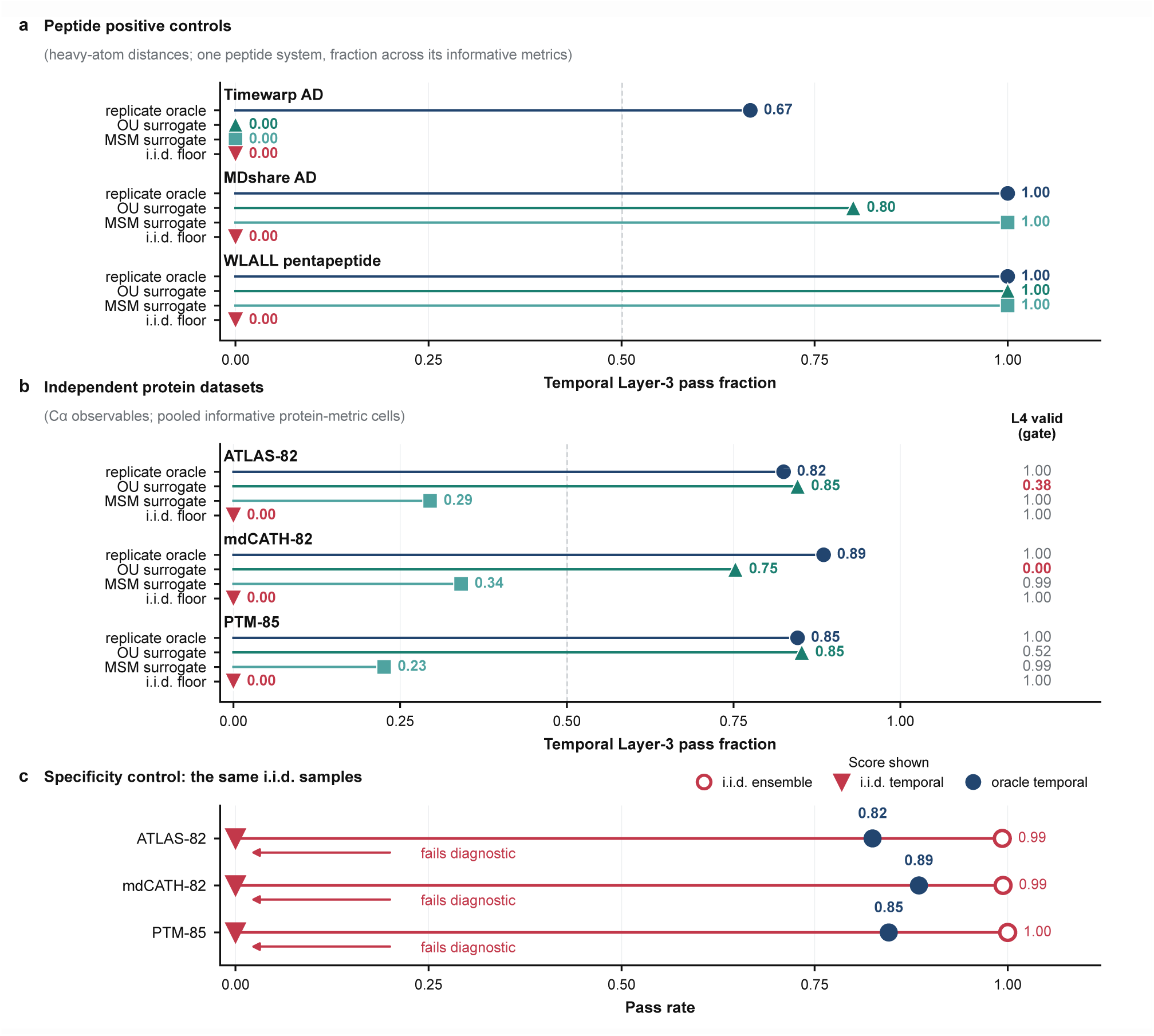
Dynbench detects reproducible temporal structure when it is present. Panels show the fraction of informative time-sensitive tests passed before geometric gating; dashed lines mark the temporal-majority threshold. (a) Positive controls on two alanine-dipeptide datasets and a WLALL pentapeptide dataset. Held-out trajectories pass while shuffled controls fail. (b) The same contrast on ATLAS-82, mdCATH-82 and PTM-85; geometric-validity fractions are shown at right. The reference retains high temporal agreement and remains geometrically valid. (c) Specificity control. Independent equilibrium samples pass ensemble tests on every protein dataset but fail the temporal test, whereas held-out trajectories retain both signals. Peptide and protein panels use structural observables matched to each system.

### Submission contract and adapters

A submission is a per-protein NumPy archive with Cα coordinates (T,L,3) in reference residue order, an ordered flag and, for an ordered trajectory, a physical frame interval. Adapters may index a model trajectory by physical time, by declared frame index when no physical claim is made, or as unordered; only the physical-time branch is eligible for temporal metrics. The time-axis registry records the source interval, production stride and resulting physical interval for every ordered row. Scoring reports record aligned model and reference frame counts per protein; a matrix audit verifies every protein against the declared contract before temporal interpretation. Malformed arrays, residue-count mismatches and non-finite coordinates are rejected as contract errors. All contract-valid batches, including those flagged for collapse or blow-up by the radius-of-gyration diagnostic, proceed to scoring; only independently established pipeline faults are excluded. Reference adapters preserve each model’s declared time semantics.

### Four layers and the verdict

Layer 1 (frame realism) is a single-frame sanity check, saturated on equilibrium MD frames and not expected to rank plausible generators. Layer 2 (ensemble) compares free-energy-surface and contact-distribution divergences and a per-mode displacement Wasserstein distance^28^ against the reference ensemble. Layer 3 (kinetics), evaluated only for ordered submissions, compares time-lagged autocorrelation, VAMP-2 score^29^, implied-timescale, transition-matrix and directional-predictability statistics. Temporal metrics are evaluated only when a model supplies a physical frame interval that supports the requested lag; unsupported combinations are NA rather than interpreted as frame-index dynamics. Layer 4 tests steric clashes, virtual Cα–Cα bond-length violations, Ramachandran outliers^30^ when backbone atoms are supplied, late-versus-early drift for ordered trajectories and root-mean-square-fluctuation spread. The steric statistic is the number of non-adjacent Cα pairs below 4 Å divided by all eligible frame-pairs. This pair-normalised rate removes the strong chain-length dependence of the former frame-level “any clash” fraction. Its absolute ceiling, 0.000758, was calibrated before rescoring as twice the maximum across 246 reference replicates; each protein also retains its reference-replicate maximum inflated by 50%, and the larger bound is used. Bond, Ramachandran and drift guardrails likewise combine a reference envelope with an absolute ceiling. RMSF spread is two-sided, failing below 0.5× the minimum or above 2.0× the maximum reference value. Radius of gyration remains a QC diagnostic and never enters the verdict. Local geometry takes precedence, followed by excessive drift or fluctuation, insufficient fluctuation and VALID.

Markov-state modelling distinguishes equilibrium agreement from kinetic validity through Chapman–Kolmogorov tests, implied-timescale convergence and related diagnostics. Dynbench operationalises this principle for heterogeneous external generators through three submission-level controls. Each metric is bracketed by target-removed and reference-derived controls, applicability is enforced by task and temporal support, and kinetic credit is withheld when a trajectory fails a separately defined geometric-admissibility gate.

Layer-4 failures enter the board by one model-agnostic rule, applied before post-gate kinetic credit. Every contract-valid ordered submission retains its ensemble and geometric readouts irrespective of Layer-4 status, and a protein passes the post-gate calculation only when it satisfies both the temporal majority and Layer 4. ProTDyn passes Layer 4 on 3 of 74 proteins; its raw post-gate rate is 0.01, but the formal post-gate kinetic pass is NR because the 10-ns contract is range-limited (Supplementary Note 2). Malformed arrays, residue-order mismatches and non-finite coordinates fail fast before scoring and are not treated as model failures.

For each metric on each protein we compute three scores: the model, a trivial surrogate (time-shuffled or i.i.d., target property removed) and a class-specific operational reference whose identity depends on the metric’s class (below). Writing T, M, O and S for the trivial surrogate, the model, the operational reference and the replicate noise floor, each metric on each protein is classified informative or SATURATED (excluded) as follows. S is the irreducible agreement of one real replicate with another, pooled over the fit replicates as a worst-case bound. It sets the SATURATED test: a metric is informative only when the trivial surrogate is meaningfully worse than S, so there is reproducible cross-replicate signal to detect. O asks whether the selected operational reference closes the T→S gap: a metric for which it does not is DATA-LIMITED under that reference and sampling. O is not a theoretical performance bound. For most kinetic metrics it is a per-mode near-harmonic fit (below), which a richer model can in principle exceed, so a MODEL-OK verdict means the model matches or beats this reference, not that it has reached an upper limit. The unqualified word oracle in this paper always denotes the held-out real-MD replicate (`classical_md`), never O. Critically, T and O are chosen per metric class, not once for the whole protocol (Fig. 2c), because a single ruler would systematically mislabel one class or the other:

All four quantities are first expressed as errors, so lower is better; a metric whose native direction is higher-is-better is negated. Define

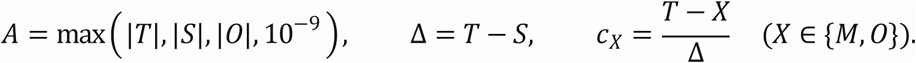

The protocol then assigns the per-metric verdict in the following fixed order:

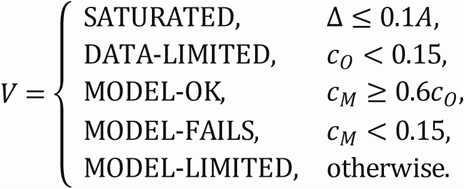

The order is part of the definition: each line is evaluated only if the preceding conditions fail. The raw closures determine the verdict; clipping them to [−5, 5] is used only for reporting.

- Equilibrium metrics (Layers 1–2). T is a static structure and O is an i.i.d. equilibrium resample of the reference. Reproducing the equilibrium marginal is the target of this class, so the resample spans the target distribution more appropriately than the kinetic OU comparator.
- Kinetic metrics (most of Layer 3). T is a time-shuffled surrogate (the equilibrium marginal with the temporal order destroyed), and O is a per-mode Ornstein–Uhlenbeck operational reference fitted on the reference trajectory, which asks whether a simple near-harmonic kinetic model suffices. It is a comparator, not an upper bound.
- Transition and mean-first-passage metrics (Layer 3). T is again the time-shuffled surrogate, but O is a held-out real replicate, because a single-basin OU process structurally cannot represent multi-state transitions and would read DATA-LIMITED on precisely the proteins whose kinetics are most interesting. Two of the seven time-ordered metrics (transition-matrix and mean-first-passage) take this replicate reference; the remaining five take the OU comparator.

The held-out MD row is scored as an ordinary submission and differs from the per-metric operational reference O. The informative set is selected once per protein and shared by the trivial floor, surrogates and deep models. Selection requires real-replicate agreement to exceed the trivial surrogate and can favour the oracle’s absolute score, while every row remains evaluated on the same selected metrics.

The leave-one-replicate-out analysis reselects the informative set and operational reference from each held-out replicate and gives oracle rates of 0.951–0.976. A fully decoupled control uses distinct replicates for selection, reference prediction and evaluation, returning 0.963 versus 0.951 for the coupled calculation (Δ +0.012; i.i.d. floor 0.00).

On informative metrics the model’s position between the trivial floor and the class-specific operational reference gives a verdict, assigned by a fixed ladder: SATURATED, then DATA-LIMITED, then MODEL-OK, then MODEL-FAILS, else MODEL-LIMITED. Reversing the MODEL-OK and MODEL-FAILS checks changed no metric cell or kinetic-pass row. Layer scores are the fraction of MODEL-OK among informative, non-SATURATED instances. MODEL-OK is the only pass: SATURATED instances leave the denominator, whereas DATA-LIMITED and MODEL-LIMITED remain and count against the model. MODEL-LIMITED accounts for 8.9% of scored cells (1,227 of 13,791).

The census runs over 13,791 scored metric-by-protein cells; a further 528 are not applicable by task, giving a 14,319-cell grid. DATA-LIMITED, the verdict that says the operational reference does not resolve the target under this sampling, accounts for 267 scored cells (1.9%) and 267 of 8,998 non-saturated cells (3.0%). SATURATED accounts for 3,762 of the 12,760 scored cells outside the Layer-1 sanity check (29.5%). On these cells the trivial surrogate already reaches the replicate noise floor, so the metric cannot discriminate submissions on that protein. The non-discriminating share varies among rows because the deep models are evaluated at different native time intervals; it is therefore reported rather than treated as model-invariant. This census quantifies which cells can support a model comparison and prevents saturated observables from diluting the layer scores.

In the public data schema, the post-gate kinetic pass rate is stored under the legacy field name `realdyn`. This rate counts a protein only if the model passes at least half of its informative temporal Layer-3 metrics and is Layer-4 valid. It measures success under the audited contract rather than a fraction of physical dynamics learned. Formally, let ℐ*_p_* be the non-SATURATED, non-NA temporal metrics for protein *p*, let *q_p_* be their MODEL-OK fraction and let *g_p_* indicate a VALID Layer-4 status:

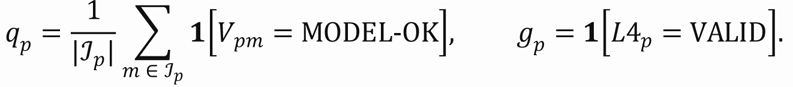

For *P*\* = {*p*: |ℐ*_p_*| > 0},

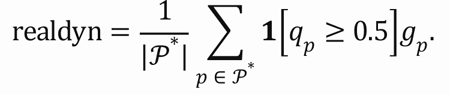

Unordered ensemble submissions have no temporal metrics by contract, so they do not enter *P*\* and the post-gate kinetic pass is NA by task rather than zero. The kinetics column of Table 1 and the pre-gate marker of Fig. 3b are the same temporal-only score over the seven time-ordered metrics used by the gate. A separate legacy diagnostic adds an eighth, order-invariant per-mode displacement Wasserstein distance. The temporal-only and legacy diagnostics differ by up to 0.09; Table 1 uses the temporal-only definition and reports ensemble submissions as NA.

For the learned-output synthetic-order stress test, we selected the first 50 available frames after the frozen model-specific preprocessing for BioEmu, P2DFlow and Str2Str, then applied ten independently seeded random permutations. One BioEmu protein with fewer than 50 frames was excluded, leaving 81, 65 and 82 proteins, respectively. Direct coordinate comparison verified that every seed used the same selected frames and changed only their order; no selected model frame matched the held-out reference coordinate fingerprint. The permuted outputs were scored as ordered submissions solely to probe order-sensitive metrics. This within-ensemble stress test leaves production task labels and scorecard entries unchanged (Supplementary Note 9).

The kinetic-pass denominator contains all non-SATURATED temporal tasks; a DATA-LIMITED cell remains a non-pass because it reflects unresolved control or data rather than model failure. Excluding these cells raises the oracle from 0.951 to 0.976, ConfRover from 0.247 to 0.288 and the MSM comparator from 0.317 to 0.341, while MDGen remains 0.134 and the i.i.d. floor 0.00. Table 1 uses the full denominator.

Temporal support depends on the submitted time interval. We therefore require a contract-specific reference-support audit before reporting a post-gate kinetic pass: the oracle-minus-floor separation must be at least 0.30 with a 95% bootstrap interval excluding zero, and the median number of informative temporal metrics must be at least three. These post hoc operational criteria were fixed before the leak-free 105-cell audit. ConfRover and MDGen pass at 1 ns, with median support four and separations 0.956 (95% CI, 0.903–0.996) and 0.921 (0.859–0.971). BioKinema and ProTDyn retain three metrics at 10 ns but are range-limited, with separations 0.053 (0.032–0.076) and 0.058 (0.036–0.085). MarS-FM is support-limited at approximately 50 ns, with three reference frames, median support one and separation 0.034 (0.018–0.056). Excluding low-resolution MSM metrics does not change any status. The audit uses 20 independent i.i.d. seeds and 2,000 hierarchical bootstrap draws. A 5-by-3 sensitivity grid spanning gap thresholds 0.20–0.40 and median-support thresholds 2–4 retains ConfRover and MDGen as resolved and the other three contracts as unresolved in all 15 settings. Because the contract criterion uses median support whereas the post-gate denominator admits any protein with at least one informative metric, a separate sensitivity excludes proteins below per-protein minima of two, three or four before protein bootstrap.

Reference-only temporal-resolution curve. We repeated the leak-free reference-support audit on all 82 ATLAS proteins over 12 conditions. A fixed 100-ns horizon used intervals of 1, 2, 4, 5, 10, 20 and 50 ns, retaining 101, 51, 26, 21, 11, 6 and 3 frames. A fixed 11-frame arm used intervals of 1, 2, 4, 5 and 10 ns, spanning horizons of 10–100 ns. Replica C remained the evaluation target, replicas A and B formed the fit and i.i.d.-floor pool, and B served as the reference stand-in. Each condition used 20 independently seeded floors and 2,000 hierarchical bootstrap draws. The same gap and informative-support criteria defined reportability. This post hoc grid maps the tested ATLAS reference and contains no learned-model score (Extended Data Fig. 5b and Supplementary Note 10).

Stationary-matched transition-rate perturbation control. For each ATLAS protein, we fitted a reversible microstate Markov model to replicas A and B on four reference-fit TICA modes, with a PCA fallback, and retained replica C for evaluation. From the one-step matrix P we formed operator arms 0.75I + 0.25P, 0.5I + 0.5P, P, P² and P⁴, labelled by nominal dynamical-speed factors of 0.25, 0.5, 1, 2 and 4. Stationary state counts were rounded once per protein and seed, authentic fit-replicate frames were sampled to those counts and the exact same coordinate multiset was reordered under every arm. Thus the five conditions differed in microstate order but not coordinates or populations. Five fixed seeds were averaged within protein before 2,000 protein-bootstrap draws. Primary contrasts were the 0.25-fold and 4-fold arms versus the fitted chain; Layer-2 and Layer-4 point changes no larger than 0.05 defined marginal and admissibility stability. The labels index transition-operator perturbations rather than exact scaling of a continuous-time physical generator (Extended Data Fig. 7 and Supplementary Note 12).

The post-gate kinetic pass is a stringent per-protein binary rate, and its graded companion is the temporal Layer-3 informative-pass rate. Together they separate the fraction of proteins satisfying the post-gate criterion from the average fraction of informative temporal tests passed. Both require a resolved contract-level reference-support audit.

### Baselines and anchors

Four baselines instantiate the ladder, each scored as a submission.classical_md is a held-out real-MD replicate and serves as the oracle anchor. msm_generative is sampled from a Markov state model^25^, which represents hopping among discrete conformational states fitted in the reference TICA^31^ coordinates.tica_ou is a per-mode Ornstein–Uhlenbeck / Koopman surrogate, and iid_equilibrium is an i.i.d. resample of reference frames. The resulting ladder follows the expected order (Table 1): classical MD is the strongest post-gate anchor, the MSM reproduces ensemble and transition statistics, the i.i.d. resample passes ensemble but fails kinetics, and the OU surrogate passes near-harmonic kinetics but not full validity.

### Sensitivity analyses

Six analyses test sensitivity to subset composition, sequence overlap, geometric guardrails, temporal thresholds, operational-reference choice and reference-assisted calibration.

Subset bias. The five ordered deep models and four kinetic anchors share 72 proteins. Recomputing every row on these same protein IDs gives oracle post-gate kinetic pass 0.972, MSM 0.361, OU 0.319, ConfRover 0.250 and MDGen 0.083. The same calculation retains BioKinema 0.069, ProTDyn 0.014 and MarS-FM 0.000 only as raw diagnostics because their contracts remain unresolved; the i.i.d. floor is 0.000.

Sequence overlap. Two screens are reported. The frozen protocol used exact sequence matching and k-mer-Jaccard overlap (k = 3, Jaccard ≥ 0.3) against a combined ATLAS training profile, flagging 4 of 82 proteins. We then ran a post hoc alignment-based audit because identity alone admitted short local matches. MMseqs2 at sensitivity 7.5 searched the 82 evaluation chains against 1,266 ATLAS training chains. A protein is flagged when a hit reaches at least 30% sequence identity and at least 50% query coverage. The audit flags 10 proteins, leaving 72 unflagged. The five ordered deep models and four kinetic anchors share 72 proteins, of which 62 are unflagged. On these same proteins the screened-minus-unscreened paired differences are small and every 95% bootstrap interval includes zero; the oracle-to-floor separation changes by −0.004 (95% CI, −0.013 to 0.000). An unflagged protein has no above-threshold match in the auditable corpus.

Geometric-guardrail calibration. Each Layer-4 guardrail fires when a submission exceeds both a fixed absolute ceiling and its protein-specific reference-replicate envelope inflated by a margin (Four layers and the verdict). Under this rule, the held-out real-MD replicate is valid on 82 of 82 proteins. Threshold sensitivity is reported below. Under the pair-normalised clash statistic, ConfRover is invalid on 16 of 73 proteins and BioEmu on 24 of 82.

Threshold robustness. The post-gate kinetic pass applies a 0.5 pass-fraction threshold to a protein’s informative Layer-3 metrics. Swept over {0.3, 0.4, 0.5, 0.6, 0.7}, the oracle stays 0.79–0.99 and the i.i.d. floor remains 0.00. Intermediate values are sensitive: ConfRover ranges from 0.40 to 0.03 and MDGen from 0.13 to 0.06; the unresolved BioKinema raw diagnostic ranges from 0.24 to 0.00. The anchor bracket is stable, while deep-model ordering remains threshold-dependent (Extended Data Fig. 4a).

Oracle/floor stability. The threshold sweep and leave-one-protein-out jackknife cover the two anchors, two surrogates and five ordered deep models. No jackknife row moves by more than 0.014 (oracle 0.012, ConfRover 0.014 and MDGen 0.012; the unresolved ProTDyn and BioKinema raw diagnostics move by 0.014 and 0.013, respectively; the i.i.d. floor and MarS-FM remain 0.000). The anchor bracket is therefore stable to the removal of any one protein (Extended Data Fig. 4b,d).

Operational-reference sensitivity. In the frozen protocol, most temporal relaxation metrics use a fitted Ornstein-Uhlenbeck process as their operational reference, while transition and mean-first-passage metrics use a held-out replicate. A post hoc sensitivity reroutes all seven genuinely temporal metrics to the held-out replicate while retaining the time-destroyed and pooled-replicate floors. Temporal raw quantities were required to match the frozen reports before verdict replacement; non-temporal metrics and Layer 4 were unchanged. Protein-paired intervals use 2,000 bootstrap resamples. The same analysis was applied to the completed 4-ns ConfRover mdCATH rollout.

Reference-assisted calibration diagnostic (S1c). S1c tests whether part of a model’s post-gate deficit reflects Layer-4 rejection of trajectories that already pass the temporal criterion. It is applied to ConfRover and MDGen, the two ordered contracts that cleared the reference-support gate before this analysis. Each protein supplies three replicates: the last is the evaluation target, one is the N1 reference stand-in and the remaining replicate supplies calibration statistics. The evaluation target never enters calibration or the N2 pool. Fluctuations are rescaled about the mean structure to match the calibration-replicate root-mean-square fluctuation, and virtual Cα–Cα bonds are projected onto the reference bond length. The scale is estimated on the 100-frame aligned view used by Layer 3 and then applied to the complete native rollout. Temporal metrics use the same 100-frame window, whereas Layer 4 is recomputed on the corrected complete rollout. The correction is applied symmetrically to model, N1 and N2 arms. Attainment is the corrected model rate minus the corrected N2 rate divided by the corrected N1-minus-N2 range. The N2 floor uses 20 seeds whose coordinates and sampling indices are reused from the uncorrected run. Confidence intervals use 2,000 bootstrap draws resampling proteins and seeds independently. The post hoc between-model interaction uses the 73 shared proteins. S1c requires same-protein reference MD and therefore remains a diagnostic of gate effects (Extended Data Fig. 3).

Collapse and blow-up quality control. The submission writer records the median radius-of-gyration ratio and flags values below 0.5× or above 2× the reference. Contract-valid coordinates continue to scoring, where Layer-4 status is determined by virtual-bond, clash, drift and fluctuation guardrails; radius of gyration remains a separate diagnostic. Only independently established pipeline faults, such as an adapter, unit or checkpoint mismatch, trigger exclusion, leaving genuine collapse and blow-up visible as model outcomes.

### Uncertainty, matched subsets and versioning

Scorecard intervals use 1,000 protein-cluster bootstrap resamples (seed 0). Layer-2 and temporal Layer-3 scores are pooled MODEL-OK fractions over informative protein-metric cells, with proteins resampled as clusters; Layer-4 validity and reportable kinetic-pass values are per-protein fractions. Partial coverage is not imputed. The five ordered deep models and four kinetic anchors are additionally recomputed on their 72-protein-ID intersection, while P2DFlow is reported on its own 65-protein coverage and enters no ordered comparison. A leave-one-protein-out jackknife moves no raw post-gate row by more than 0.014. Repeating the oracle calculation with each MD replicate held out gives 0.976, 0.976 and 0.951; the i.i.d. floor remains 0.00.

The verdict-constant sweep is interpreted at the anchor level. Across 27 combinations of saturation tolerance, learnability tolerance and MODEL-OK ratio, the oracle-to-floor separation remains 0.80–1.00 and the i.i.d. floor remains 0.00. Intermediate deep-model values and ordering are not invariant: at the strictest MODEL-OK ratio ConfRover and MDGen can exchange order, and BioKinema reaches the floor under the strictest kinetic-majority threshold. The Layer-4 values use pair-normalised clash counts. A Layer-4 guardrail sweep replays the stored per-protein guardrail quantities across the tunable constants; at the frozen defaults the replay reproduces all twelve Layer-4 rows exactly. The anchor separation is invariant: the held-out MD anchor holds 1.000 and MarS-FM 0.000 at every setting. Widening or narrowing the clash envelope changes almost nothing, and rescaling the calibrated clash ceiling from 0.25× to 4× shifts absolute values (MDGen 0.000–0.171, ConfRover 0.658–0.836) without reordering the five ordered rows. One constant is genuinely load-bearing: as the RMSF too-cold fraction rises from 0.3 to 0.7, ConfRover falls from 0.849 to 0.452, so its majority Layer-4 validity survives a fraction of 0.6 but not 0.7, whereas BioKinema stays above 0.92 throughout (Supplementary Note 6).

### Statistics and reproducibility

The unit of resampling is the protein. Pooled Layer-2 and temporal Layer-3 fractions use a protein-cluster bootstrap; Layer-4 and kinetic-pass values use a protein-level bootstrap (1,000 resamples, seed 0). The Fig. 1d temporal ablation and sequence-overlap sensitivity use 2,000 protein-level resamples. Binary fractions use Wilson or bootstrap intervals as stated in each legend. No statistical test predetermined sample size; the evaluation set is the full 82-protein split. Not-applicable-by-task cells are excluded rather than scored zero, and partial coverage is not imputed. The split, anchors and verdict rules were frozen before model promotion, but the alignment audit and pair-normalised clash analysis are explicitly post hoc. This is a computational benchmark, so randomization and blinding of biological samples do not apply.

Generative AI tools (OpenAI Codex and Anthropic Claude) assisted code review, manuscript editing and consistency checks. The authors independently verified all analyses, numerical results, citations and final text and take full responsibility for the work.

### Capability matrix

Extended Data Table 1 compares each prior evaluation across seven axes: ensemble fidelity, ordered or temporal kinetics, metric-specific floor, ceiling and noise-floor calibration, geometric-admissibility gating, applicability-by-task, a sequence-overlap-screened shared set and a frozen public leaderboard. Every entry is tied to a named table, figure or section in Supplementary Note 1. Yes denotes evaluation on proteins and Partial denotes evaluation on peptides only. MDGen, MarS-FM and Timewarp are therefore marked Partial for kinetics because peptide torsions are not the same measurement as protein-domain dynamics. The geometric-gate column records whether geometry gates the dynamical verdict, not whether geometry is merely reported. Str2Str defines a validity metric class and ConfRover reports MolProbity geometry, but neither uses geometry as a gate. Beyond Ensembles reports a protein-level kinetic entry as unstable and separately defines long-horizon failure by an lDDT-based structural criterion^18^. Several methods populate the first two axes, and individual studies implement parts of the remaining protocol. None combines all five protocol-level axes added by Dynbench.

## Extended Data

**Extended Data Fig. 1.**
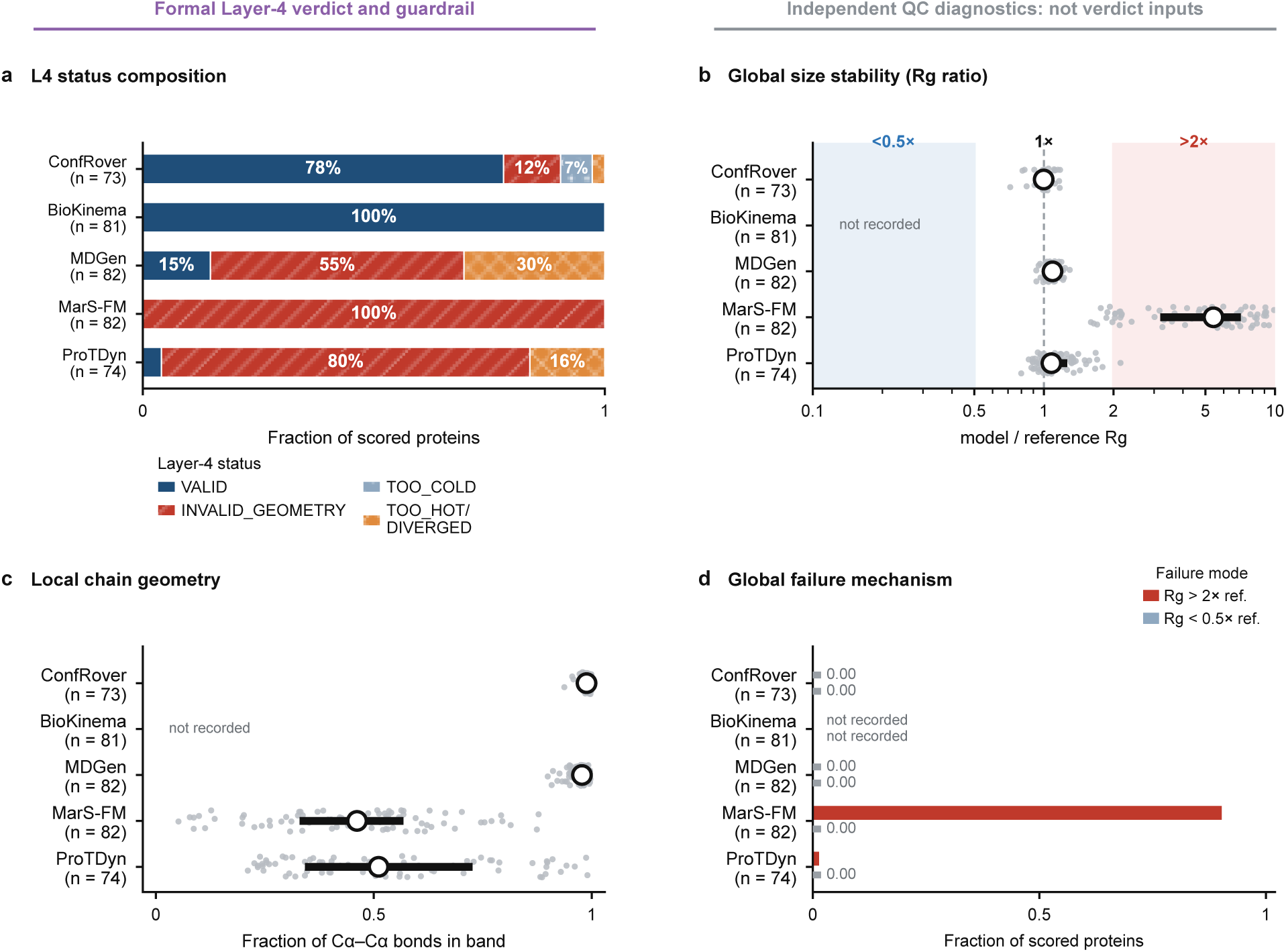
Geometric quality control across ordered models. (a) Layer-4 status after applying the pair-normalised steric-clash statistic and the other geometry and fluctuation guardrails (ConfRover, 78% valid; BioKinema, 100%; MDGen, 15%; MarS-FM, 0%; ProTDyn, 4%). (b) Per-protein radius-of-gyration ratios relative to reference MD. The shaded regions mark independent collapse and blow-up diagnostics; radius of gyration is not a Layer-4 verdict input. (c) Per-protein fractions of virtual Cα–Cα bonds within the accepted band, shown as a local-chain diagnostic. (d) Fractions of scored proteins with global size failure (radius of gyration above twofold or below one-half of the reference value). Auxiliary radius-of-gyration and bond-level distributions were not recorded for BioKinema and are marked accordingly in b–d. MarS-FM fails globally on most proteins, whereas MDGen’s Layer-4 failures arise mainly from the formal geometry guardrails rather than global expansion.

**Extended Data Fig. 2.**
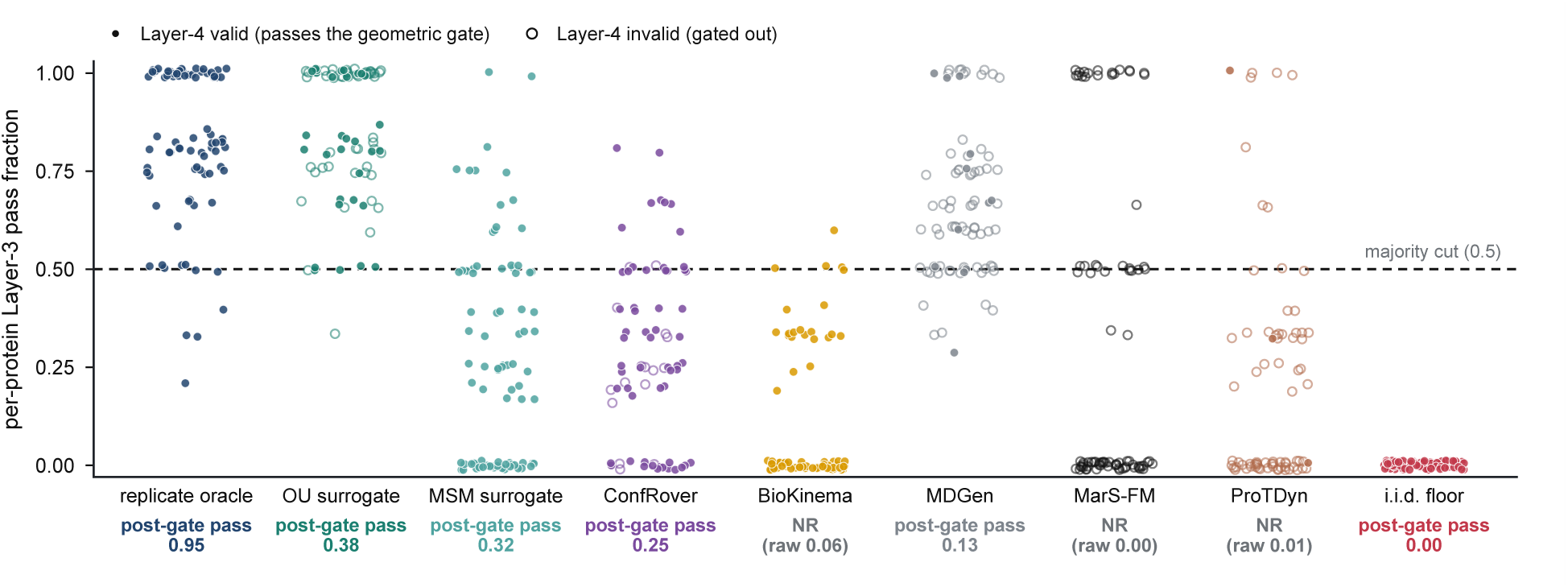
Per-protein temporal diagnostics and the Layer-4 gate. Per-protein native-task Layer-3 informative-pass fractions for models scored on kinetics; each point is one protein, filled where the trajectory is Layer-4 valid and hollow where it is invalid. For resolved contracts, post-gate kinetic pass is the fraction meeting both the 0.5 temporal-majority threshold and Layer 4. BioKinema and ProTDyn are marked NR because their 10-ns contracts are range-limited, and MarS-FM because its approximately 50-ns contract is support-limited; their raw post-gate values are printed only for provenance. MarS-FM remains invalid on every protein, whereas BioKinema is valid on every scored protein. Native output intervals differ among deep models, so the panel diagnoses within-row gate behaviour rather than a common-resolution temporal ranking.

**Extended Data Fig. 3.**
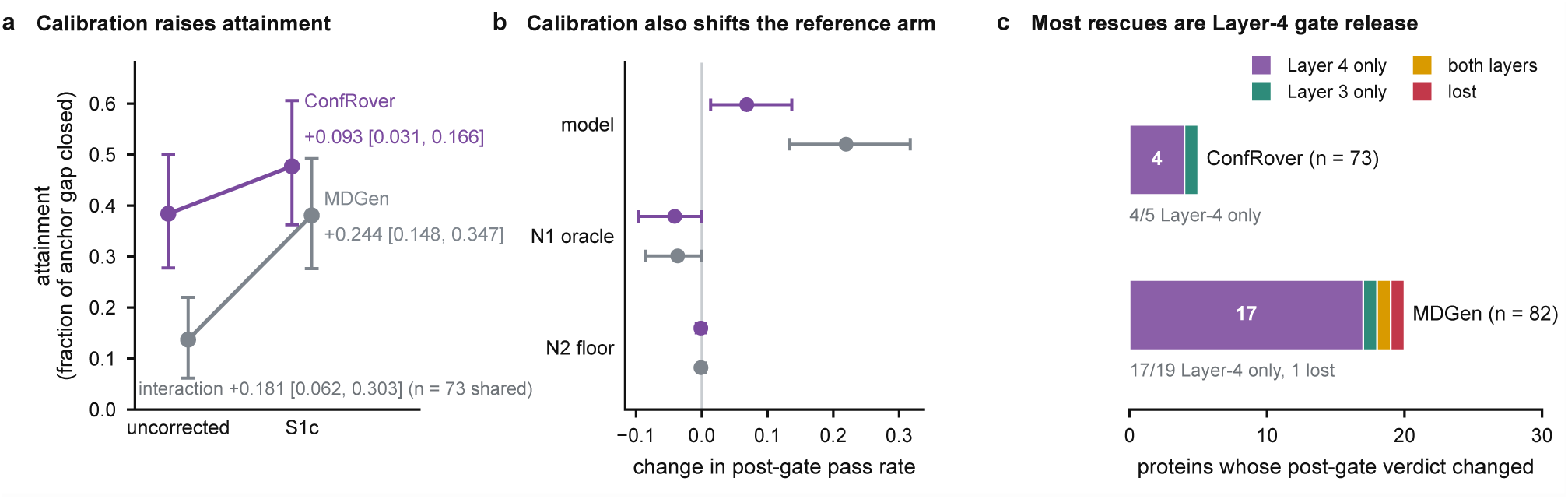
Reference-assisted calibration separates temporal and geometric failure modes. S1c rescales fluctuations to an independent same-protein MD replicate and projects virtual Cα–Cα bonds to reference lengths. It is applied symmetrically to the model, N1 reference and N2 floor under a 100-frame support-matched contract. (a) Attainment, the fraction of the empirical N1–N2 gap closed, before and after S1c. Estimates and 95% confidence intervals use 2,000 bootstrap draws over proteins and 20 N2 seeds. Paired changes are +0.093 (0.031 to 0.166) for ConfRover and +0.244 (0.148 to 0.347) for MDGen; their post hoc interaction is +0.181 (0.062 to 0.303; 73 shared proteins). (b) Changes in each arm. S1c shifts the ConfRover N1 arm by −0.041 (−0.096 to 0.000), motivating correction of all arms before ratio formation. (c) Sources of post-gate changes. Four of five ConfRover rescues and 17 of 19 MDGen rescues were Layer-4-only. Each model had one Layer-3-only rescue; MDGen also had one joint rescue and one protein that lost its temporal pass while remaining Layer-4 valid. Layer 4 was recomputed on each corrected complete trajectory. S1c requires same-protein reference MD and is therefore diagnostic.

**Extended Data Fig. 4.**
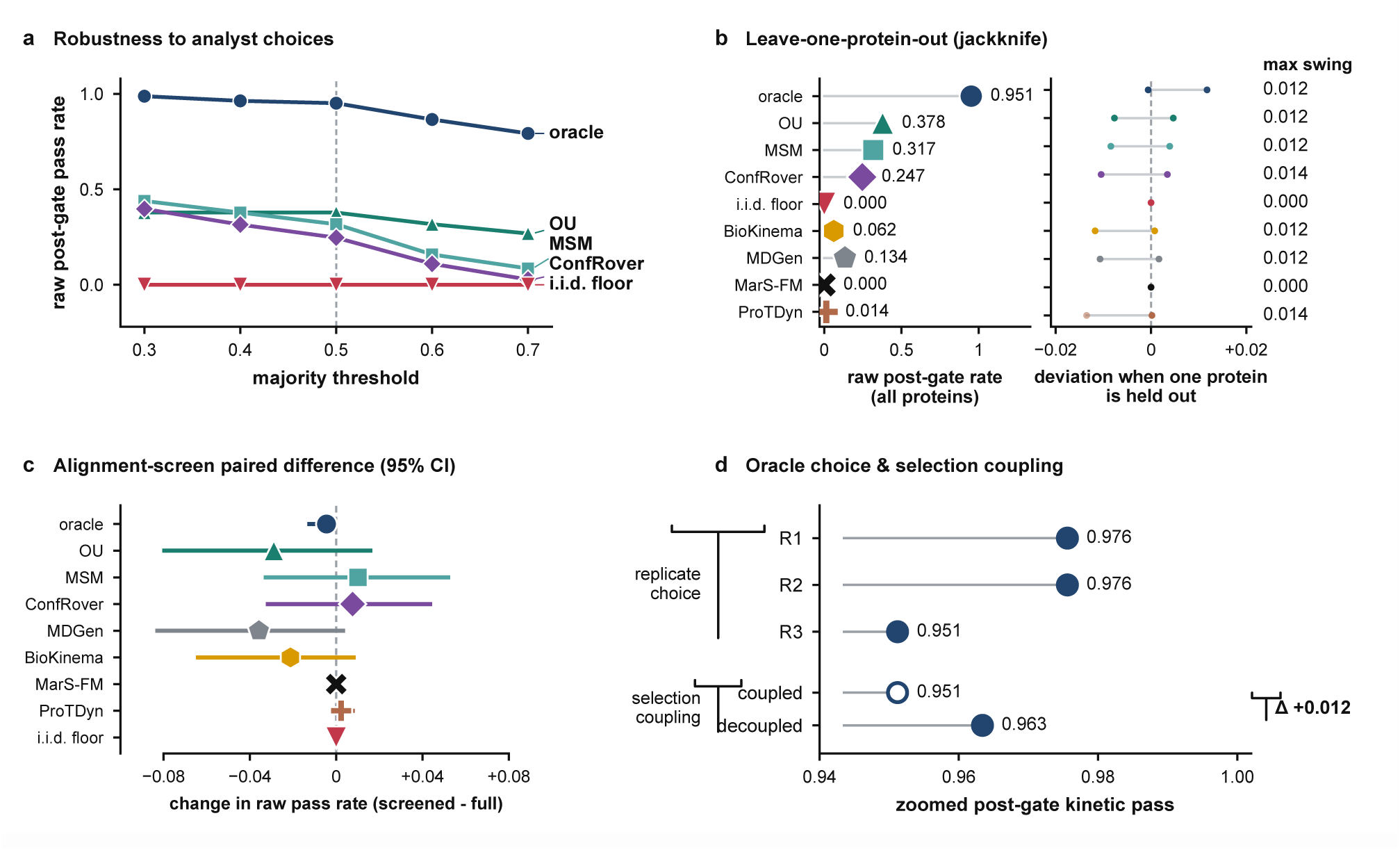
The anchor bracket is robust, whereas raw intermediate diagnostics remain threshold-dependent. (a) Post-gate kinetic pass rate versus the temporal-majority threshold for two anchors, two surrogates and five ordered deep models. The oracle remains 0.79–0.99 and the i.i.d. floor 0.00. BioKinema reaches the floor at 0.7 and ConfRover declines to 0.027. (b) Leave-one-protein-out jackknife for the same raw rows; none moves by more than 0.014. (c) Post hoc alignment-based sequence-overlap audit on the 72 shared proteins, of which 62 are unflagged. Every paired interval includes zero and the oracle-to-floor separation changes by −0.004 (95% CI, −0.013 to 0.000). (d) Holding out each real-MD replicate in turn gives oracle post-gate kinetic pass 0.951–0.976; a fully decoupled control gives Δ = +0.012.

**Extended Data Fig. 5.**
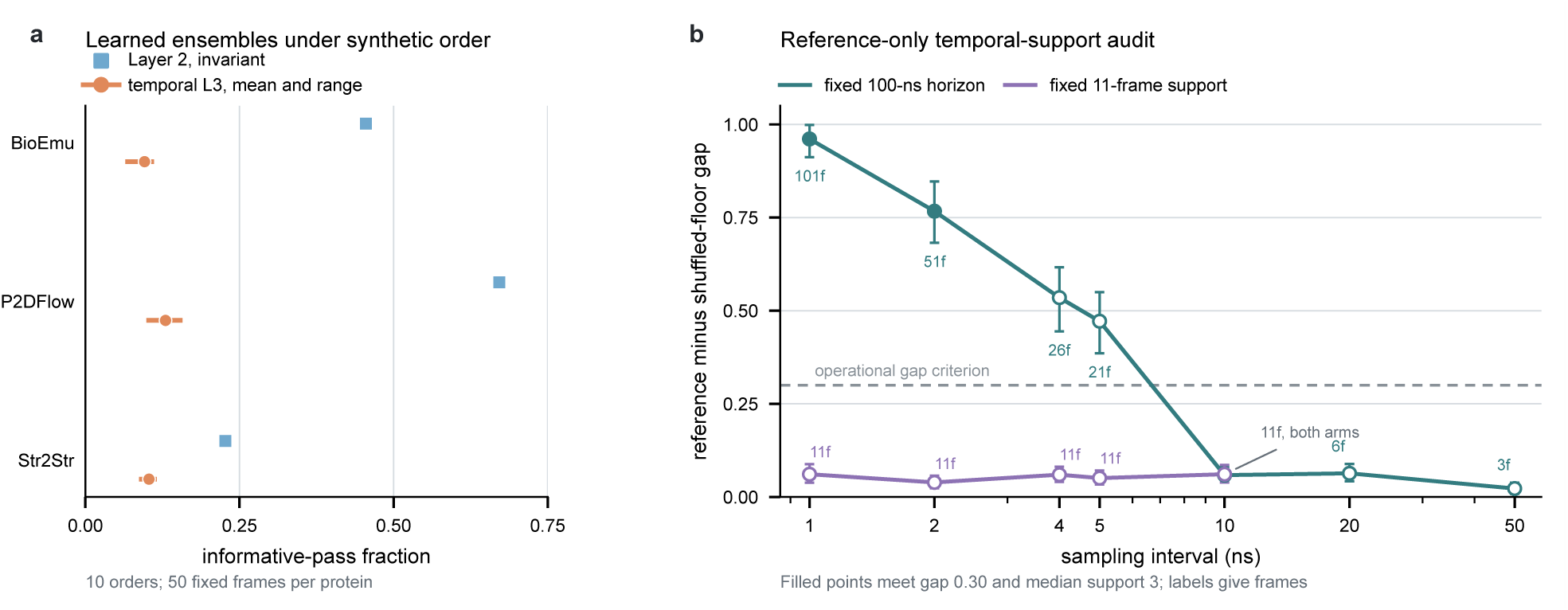
Learned ensembles retain distributional scores under arbitrary ordering, and temporal reportability requires both resolution and horizon. (a) Fixed 50-frame outputs from three unordered generators were assigned ten independently seeded synthetic orders. Squares show the invariant Layer-2 informative-pass fraction; circles and horizontal ranges show the mean and range of temporal Layer-3 fractions. Coverage was 81 proteins for BioEmu, 65 for P2DFlow and 82 for Str2Str. Direct coordinate comparison confirmed identical frame sets across orders and no held-out-reference coordinate matches. These synthetic orders probe sensitivity without supplying physical time. (b) Reference-only ATLAS audit across a fixed 100-ns horizon and fixed 11-frame support. Points show the held-out-reference minus i.i.d.-floor gap; error bars are 95% intervals from 2,000 hierarchical bootstrap draws over 82 proteins and 20 floor seeds. Filled points meet both the 0.30 gap and median-three-metric support criteria. Only the 1- and 2-ns fixed-horizon conditions are reportable. Source data are provided with this paper.

**Extended Data Fig. 6.**
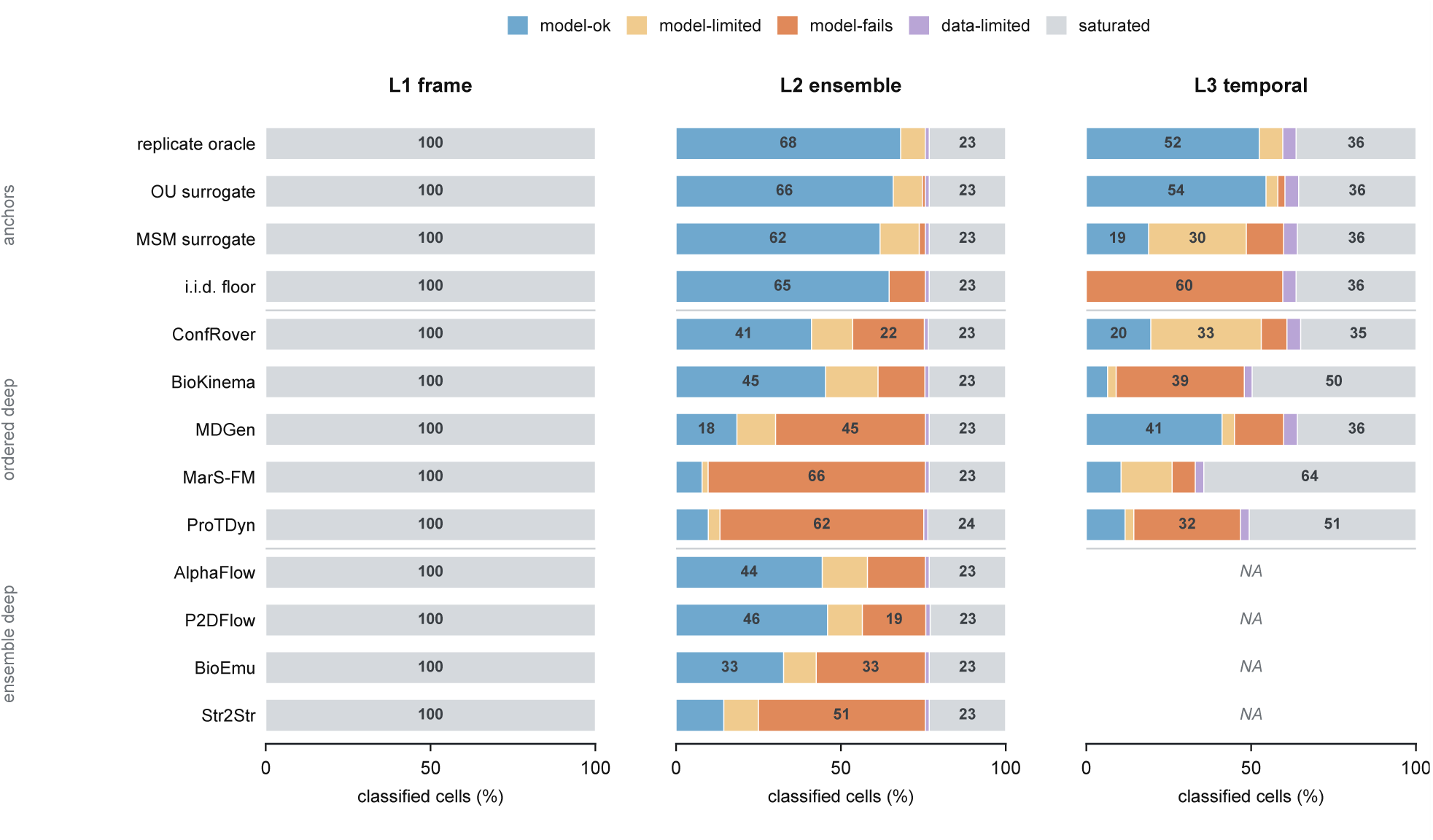
Verdicts separate by what each measurement layer tests. Bars show the five classified outcomes within Layer 1, Layer 2 and the seven time-ordered Layer-3 metrics. Each bar is normalised over all classified cells in that model and layer, so saturated and data-limited tests remain visible; MODEL-OK is the only pass. Layer 1 is saturated for every row. The i.i.d. floor is predominantly MODEL-OK in Layer 2 but MODEL-FAILS in temporal Layer 3, whereas ensemble-only models are NA by task in Layer 3. The separate order-invariant displacement diagnostic does not enter the temporal panel. Percentages are rounded and may sum to 100 ± 1. Source data are provided with this paper.

**Extended Data Fig. 7.**
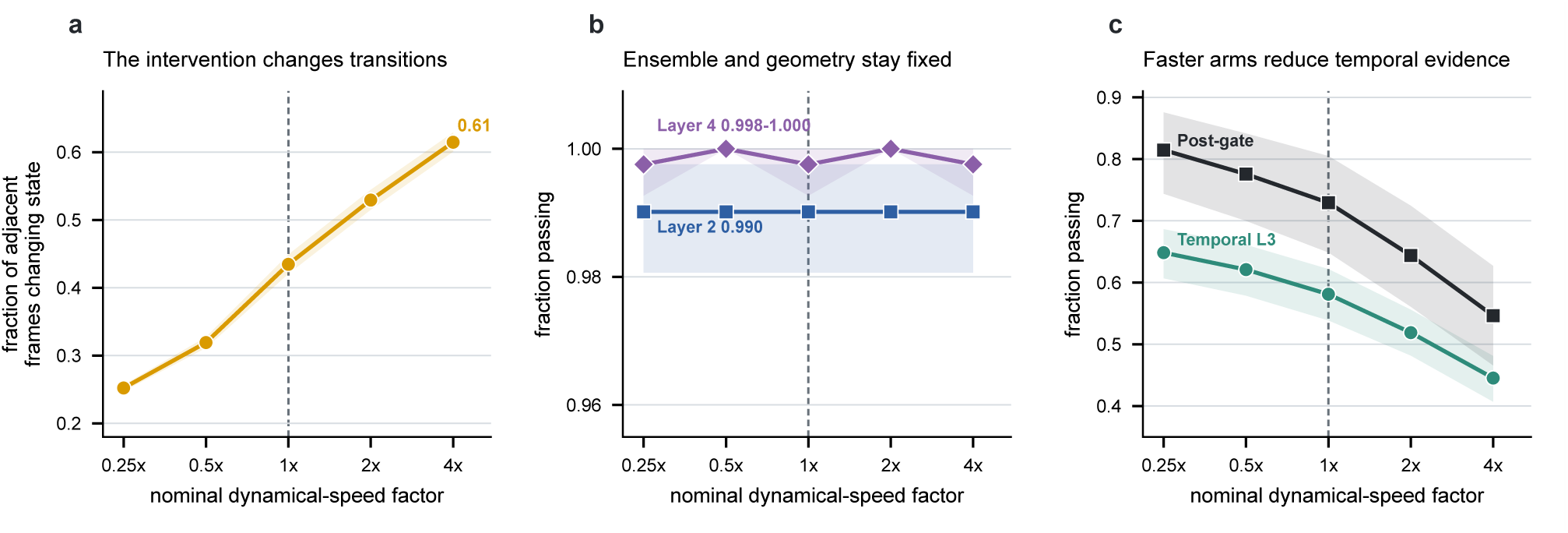
A stationary-matched control isolates transition-rate perturbation from ensemble and geometric quality. Each protein and seed uses one exact multiset of authentic fit-replicate coordinates, reordered under reversible transition operators with nominal dynamical-speed factors from 0.25-fold to 4-fold. (a) The fraction of adjacent frames changing microstate rises across the operator arms. (b) Layer-2 ensemble fidelity remains 0.990 throughout and Layer-4 validity remains 0.998–1.000. (c) Temporal Layer 3 and the post-gate kinetic pass respond monotonically to the perturbation. In the pre-specified extreme contrasts, the 0.25-fold arm changes temporal Layer 3 by +0.067 (95% CI, +0.046 to +0.090) and post-gate kinetic pass by +0.085 (+0.046 to +0.129), whereas the 4-fold arm changes them by −0.136 (−0.164 to −0.107) and −0.183 (−0.239 to −0.124). Lines show means and shading 95% protein-bootstrap intervals after averaging five fixed seeds within each of 82 proteins (2,000 resamples). The fitted chain is an operational control, not a physical rate optimum.

**Extended Data Table 1.**
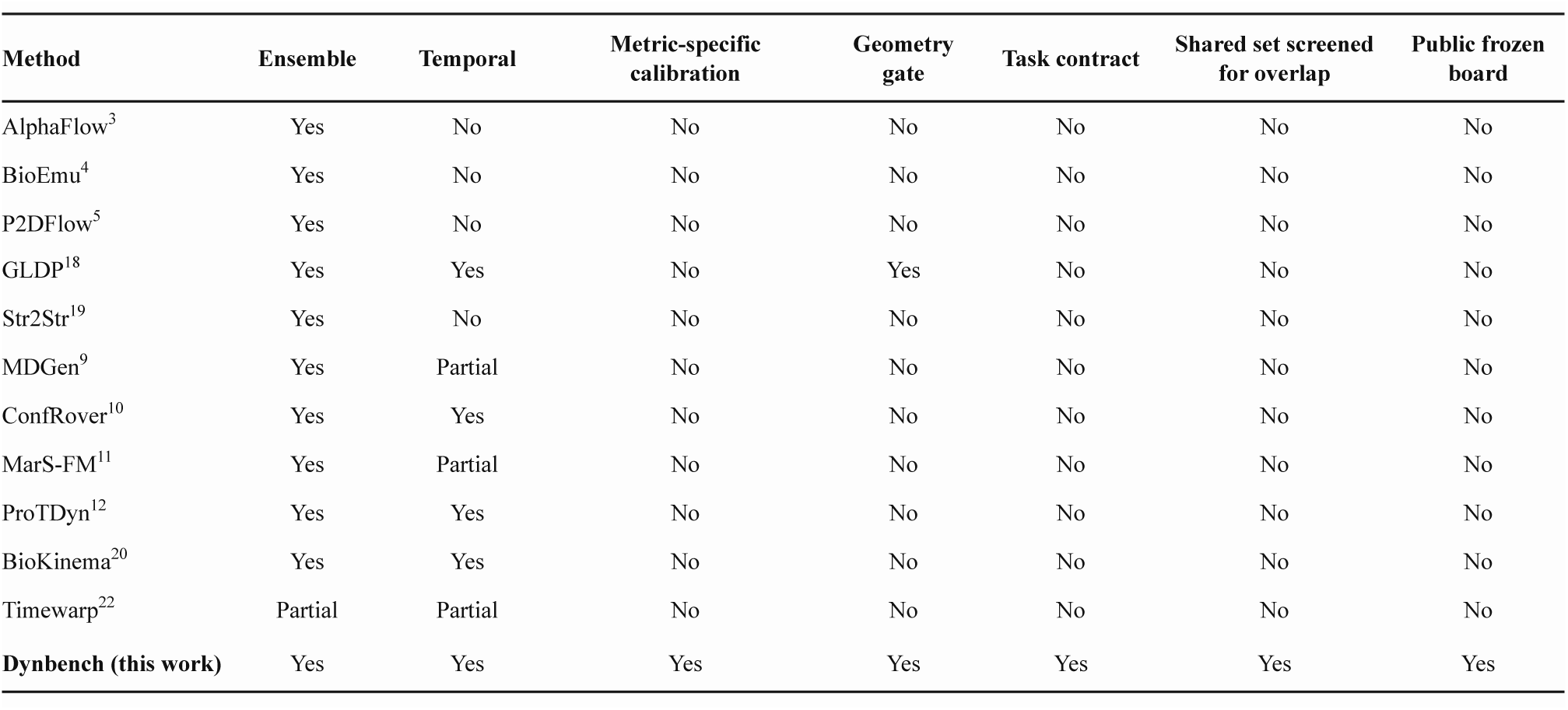
Capability matrix of prior evaluations. What each method’s original paper evaluated across the seven axes Dynbench formalises (`Yes` = evaluated on proteins; Partial = evaluated on peptides only;No = not evaluated). Every entry is justified against a named table, figure or section of the original paper in Supplementary Note 1. ProteinConformers^14^ benchmarks conformational diversity and plausibility, and Score Dynamics^16^ learns a large-timestep evolution operator on smaller systems; both are included as related methods rather than scorecard rows. Later methods are catalogued in Supplementary Table S7.

| Method | Ensemble | Temporal | Metric-specific calibration | Geometry gate | Task contract | Shared set screened for overlap | Public frozen board |
| --- | --- | --- | --- | --- | --- | --- | --- |
| AlphaFlow<sup>3</sup> | Yes | No | No | No | No | No | No |
| BioEmu<sup>4</sup> | Yes | No | No | No | No | No | No |
| P2DFlow<sup>5</sup> | Yes | No | No | No | No | No | No |
| GLDP<sup>18</sup> | Yes | Yes | No | Yes | No | No | No |
| Str2Str<sup>19</sup> | Yes | No | No | No | No | No | No |
| MDGen<sup>9</sup> | Yes | Partial | No | No | No | No | No |
| ConfRover<sup>10</sup> | Yes | Yes | No | No | No | No | No |
| MarS-FM<sup>11</sup> | Yes | Partial | No | No | No | No | No |
| ProTDyn<sup>12</sup> | Yes | Yes | No | No | No | No | No |
| BioKinema<sup>20</sup> | Yes | Yes | No | No | No | No | No |
| Timewarp<sup>22</sup> | Partial | Partial | No | No | No | No | No |
| **Dynbench (this work)** | Yes | Yes | Yes | Yes | Yes | Yes | Yes |

## Statements

### Data availability

ATLAS reference trajectories are public at https://www.dsimb.inserm.fr/ATLAS/. Dynbench 1.0.0 has been prepared under reserved DOI 10.5281/zenodo.22146996 and will be made public at publication. The archive contains the evaluation set, Cα reference bundle, submission schema, frozen ATLAS-82 scorecard, per-protein reports, quality-control cards, source data and audit records underlying the manuscript, including the frozen Lockbox-36 protocol, Broad and Deep corrected reports, technical-NA records and sensitivity analyses. mdCATH is public through its original data record. The PTM-85 supporting-control set is drawn from DynaMo-PTM, an in-house database being prepared for separate publication; until that release, it is available from the corresponding author on reasonable request. The peptide audits use the public Timewarp and MDshare alanine-dipeptide and WLALL pentapeptide datasets.

### Code availability

Dynbench scoring, model adapters, analysis scripts and leaderboard versions are publicly available under the MIT licence at https://github.com/kkkniengLiu/dynbench. The first public release is available at https://github.com/kkkniengLiu/dynbench/releases/tag/v1.0.0. Model authors may submit corrections or re-scored outputs against a named board version.

### Author contributions (CRediT)

K.L. conceived and designed the benchmark protocol, implemented the scoring pipeline and model adapters, ran all experiments and analyses, and wrote the manuscript. Q.Q. contributed to dataset preparation and model adaptation and helped revise the manuscript. Y.C. supervised the study, guided the design and analysis, and edited the manuscript. All authors read and approved the final manuscript.

