## Supplementary Information for "Ensemble tests mask missing dynamics in protein conformational generators"

**Contents.** Supplementary Notes 1-12 and Supplementary Tables S1-S8.

### Guide to the supplementary evidence

The main text presents the shortest route through the argument. This Supplementary Information provides the corresponding stress tests, implementation detail and complete aggregate results. The evidence is grouped by the question a reader may wish to test:

| Question raised by the main text | Supporting evidence | Location |
|---|---|---|
| Does the measurement distinguish an intact temporal process from the same states without their order? | Metric definitions, reference-derived anchors, threshold and replicate sensitivity, synthetic ordering of learned ensembles, and a stationary-matched transition-rate control | Supplementary Notes 4-6, 9 and 12; Tables S3-S5 |
| Is the temporal target resolvable at the stated sampling interval and observation horizon? | Per-contract reference-to-floor separation, peptide positive controls, external protein datasets and the temporal-support audit | Supplementary Notes 5, 6 and 10; Tables S4-S6 |
| Do low model results reflect temporal failure, geometric failure or an unsupported output contract? | Model-specific quality-control cards, verdict census and pre-specified failure stratification | Supplementary Notes 2, 3, 7 and 8; Tables S1-S3 and S7 |
| Does the conclusion survive a prospective dataset shift? | Frozen Lockbox-36 design, Broad and Deep analyses, overlap sensitivity and metric-family leave-one-out tests | Supplementary Note 11 |
| How does the protocol relate to prior evaluations, data provenance and sequence overlap? | Source-level evidence for the capability matrix, audited dataset provenance and the MMseqs2 threshold grid | Supplementary Note 1; Tables S6-S8 |

For a claim-centred reading, begin with Notes 4-6, continue to Notes 9-12, and then consult the model-specific Notes 2, 3, 7 and 8. Note 1 documents the literature comparison in Extended Data Table 1.

### Supplementary Note 1. Evidence for Extended Data Table 1

Extended Data Table 1 records which of seven evaluation axes each method's original paper provides. This note gives the specific table, figure or section supporting every entry.

**Conventions.**

- **Yes** means that the axis is evaluated on *proteins*.
- **Partial** means that the axis is evaluated on peptides only (2–8 residues), rather than on folded proteins.
- **No** means that the axis is not evaluated.

Four of the seven ordered generators surveyed here report protein-scale temporal evaluation: ConfRover, ProTDyn, BioKinema and Beyond Ensembles. MDGen, MarS-FM and Timewarp report temporal evaluation on peptides, but not on folded proteins in their original evaluations. Dynbench applies one shared protein-scale protocol to the eligible released models.

A separate mapping classifies 16 reported evaluation classes as order-invariant or genuinely temporal. Dynbench combines these observables with a same-frame permutation control, reference calibration, task contracts and a geometric gate.

#### Column 1. Ensemble fidelity

| method | mark | locator |
|---|---|---|
| AlphaFlow[3] | Yes | Table 1, "Evaluation on MD ensembles", reports pairwise-RMSD, RMSF, PCA-Wasserstein and contact-map divergences against ATLAS MD. Also see Table 4 (ESMFlow), Table 5 (ablations and normal-mode analysis) and Fig. 9 (RMSF). |

| method | mark | locator |
|---|---|---|
| BioEmu[4] | Yes | Equilibrium free-energy landscapes against MD; free-energy mean absolute error 0.91 kcal/mol over the converged test set (Results, Fig. 3); folding ΔG against experiment. |
| P2DFlow[5] | Yes | Table 1, "Evaluation on MD ensembles" ($\mathcal{J}$ = Jensen–Shannon divergence, $\mathcal{W}_2$ = 2-Wasserstein / RMWD); Fig. 3 (distributions of RMWD and weak-contact $\mathcal{J}$); Fig. 7 (RMSF vs MD). |
| Beyond Ensembles[18] | Yes | Table 1 evaluates protein systems 7JFL, 1R6W and A1AR with backbone and side-chain coordinate and dihedral JSDs plus time-averaged contact-map correlations. Section 4.3 and Table 2 compare equilibrium free-energy surfaces across three propagators in a fixed latent representation. |
| Str2Str[19] | Yes | Table 1 benchmarks the fast-folding proteins of Lindorff-Larsen et al.; §4.1 defines a *Validity* class and a symmetric Jensen–Shannon divergence on three MD-derived quantities. See Fig. 4 (contact maps) and Fig. 5 (TICA). |
| MDGen[9] | Yes | Table 4, "Median results on test protein ensembles (n = 82)", reports the ATLAS test proteins scored on the AlphaFlow ensemble metrics. |
| ConfRover[10] | Yes | Table 3, time-independent (ensemble) generation, reported as matching state-of-the-art ensemble generators; Table 2 uses Jensen–Shannon distance with precision/recall/F1 for conformational-state recovery. |
| MarS-FM[11] | Yes | Tables 2 and 3 report Pearson *r* for RMSD and RMSF, forward KL divergence and JSD of the radius of gyration on mdCATH proteins of up to 500 residues. Table 1 gives JSD on tetrapeptides. |
| ProTDyn[12] | Yes | Table 1, "Distributional similarity evaluation on CATH1 test dataset", reports JSD over radius of gyration, RMSD and TICA; Fig. 2 shows the free-energy surface along the top two TICA components. |
| BioKinema[20] | Yes | Fig. 2b–c evaluates ATLAS proteins through RMSF and ensemble distributions, including tICA- and PCA-based projections; the paper also reports protein conformational ensembles for the intended-MSA model. |
| Timewarp[22] | Partial | Conditional and Boltzmann distributions are compared against MD (Figs. 10–12), but only for alanine dipeptide, 2AA and 4AA peptides: "We evaluate Timewarp on small peptide systems" (§6). No protein ensemble is evaluated. |

### Column 2. Ordered or temporal kinetics

| method | mark | locator |
|---|---|---|
| AlphaFlow[3] | No | No time-dependent observable appears in the paper; the model emits an unordered ensemble and every reported metric is order-invariant. |
| BioEmu[4] | No | Markov state models appear **only** as a reweighting of the *training* data toward equilibrium (Supplementary §S.3.5.1), never as an evaluation target. What is evaluated is the equilibrium distribution. |
| P2DFlow[5] | No | Ensemble generator; no time-dependent observable is reported. |
| Beyond Ensembles[18] | Yes | Section 4.1 and Table 1 report the slowest TICA timescale and decorrelation time on the protein systems 7JFL, 1R6W and A1AR. The encoder-decoder is held fixed while the latent propagator is changed, making the temporal mechanism the controlled variable. |
| Str2Str[19] | No | The TICA plots (Fig. 5) are static projections of an unordered sample. The "kinetic clusters" are a colouring, not a measurement. The paper itself notes that such samplers "cannot reveal insights into kinetics directly" (Appendix G.1). |
| MDGen[9] | Partial | Kinetics is evaluated through torsion decorrelation times and MSM flux matrices (§4.1, Fig. 2) and autocorrelation functions (§4.3, Figs. 4–5). The paper states, "We focus on tetrapeptides as our main molecule class for evaluation" (§4), and its Limitations describe validation on peptide simulations. |
| ConfRover[10] | Yes | Table 1, Pearson correlations between conformational changes in generated and reference *trajectories*; Fig. 5, correlations of principal dynamic modes from tICA at varying lag times, computed on both reference and sampled trajectories; Table 2, recovery of conformational states over a 100 ns simulation. Evaluated on ATLAS proteins. |
| MarS-FM[11] | Partial | Fig. 7 reports torsion relaxation times for 4AA tetrapeptides; Fig. 8 reports backbone and side-chain torsion decorrelation for the same systems. The protein tables report structural observables only. |
| ProTDyn[12] | Yes | Fig. 4, autocorrelation of the top two TICA components from 0 to 800 ns lag time, evaluated on four test CATH1 **proteins**; Table 3, dynamical content. The paper presents this as a qualitative analysis on four systems. |
| BioKinema[20] | Yes | Fig. 3b reports a time-resolved adenylate-kinase open-to-closed transition, and Fig. 4 follows ordered protein conformational transitions over a 1-µs rollout. These are protein-scale temporal evaluations rather than unordered ensemble projections. |
| Timewarp[22] | Partial | Fig. 4 reports effective-sample-size speed-up for the slowest TICA component and the conditional transition density $p(x(t+\tau)\|x(t))$ (Figs. 11–12), all on AD, 2AA or 4AA peptides. |

### Columns 3–7. Additional benchmark axes

The remaining columns describe the benchmark functions that extend beyond a single model paper. Each definition is paired with the closest precedent in the surveyed literature.

**3. Metric-specific floor, operational reference and replicate-noise calibration.** Every metric is read *simultaneously* against a target-removed surrogate, a class-specific operational reference and a replicate-derived noise floor, so that a bare number becomes a classified verdict. The operational reference is an i.i.d. equilibrium resample for equilibrium metrics, an Ornstein–Uhlenbeck comparator for most kinetic metrics and a held-out real-MD replicate for transition and mean-first-passage metrics. It is not a theoretical upper bound, and a richer model can exceed it. No prior paper reports this complete construction. Several report one component: AlphaFlow, P2DFlow and MDGen compare against other *models* (MDGen's Table 4 reproduces AlphaFlow's numbers), which calibrates a method against its competitors but not against what the data can resolve. None reports a replicate-derived noise floor that separates model failure from a saturated metric or a target unresolved by the class-specific reference.

**4. Geometric and fluctuation admissibility gate.** Str2Str defines a *Validity* metric class (§4.1), and ConfRover reports MolProbity geometry and coarse-grained energy profiles (Table 4). BioKinema reports MolProbity and physical-stability checks in Fig. 2a. In all three cases geometry is reported **alongside** the dynamical or ensemble result, never **as a condition on the kinetic verdict**. Beyond Ensembles is a distinct partial precedent: Section 4.2 and Fig. 2 define long-horizon failure by an lDDT threshold calibrated against unphysical degradation, and Table 1 withholds a kinetic value as unstable in one protein-level comparison. It does not combine that stability check with metric-specific reference calibration, task applicability, a shared overlap-screened set or a frozen board. The broader Layer-4 gate here catches a distinct failure: MarS-FM has a raw native-task temporal Layer-3 diagnostic of 0.29 with 0/82 Layer-4-valid trajectories. Its raw post-gate value is 0.00, but the formal post-gate kinetic pass is NR because the approximately 50-ns contract is support-limited. MDGen has a higher native-task temporal Layer-3 score (0.64) but only 12/82 valid trajectories.

**5. Applicability-by-task contract.** A formal NA-by-task rule under which an ensemble-only model is scored on ensemble and validity and marked not applicable rather than failed or zero on kinetics. Prior papers do not place ensemble and ordered models on one axis at all, so the question never arises for them. It arises for a benchmark, and answering it with a zero would be a category error: it would silently rank an honest ensemble generator below a trajectory generator that produces broken output.

**6. Shared, sequence-overlap-screened cross-model evaluation set.** One evaluation split screened for training overlap and scored across *all* compared models. Each prior paper defines its own: AlphaFlow and MDGen use ATLAS splits, MarS-FM uses mdCATH with sequence-dissimilarity filtering (a leakage control for *its own* training set, not a shared evaluation set), Str2Str uses fast-folding proteins, Timewarp uses peptides. No two are the same set, so the published numbers are not mutually comparable. This is the reason a common frozen set is needed at all.

**7. Frozen public scorecard.** A versioned, publicly released scorecard with a fixed evaluation split, per-protein records and confidence intervals, to which new submissions can be added on the same ruler. No prior paper releases one.

---

#### Evidence sources

Sources: AlphaFlow (arXiv:2402.04845), BioEmu (bioRxiv 2024.12.05.626885; published in *Science*, 2025), P2DFlow (arXiv:2411.17196), Beyond Ensembles (official ICLR 2026 proceedings paper), Str2Str (arXiv:2306.03117), MDGen (arXiv:2409.17808), ConfRover (arXiv:2505.17478), MarS-FM (arXiv:2509.24779), ProTDyn (arXiv:2510.00013), Timewarp (arXiv:2302.01170). Locators refer to the numbering in those versions; where a paper has both a preprint and a journal version, the preprint numbering is used.

---

### Supplementary Note 2. ProTDyn as a QC-failing, range-limited diagnostic row

ProTDyn-dynamics covers 74 proteins, of which 59 are INVALID_GEOMETRY and 12 are TOO_HOT_OR_DIVERGED, leaving 3 Layer-4-valid trajectories. Its raw 10-ns diagnostics are temporal Layer 3 = 0.238 and post-gate kinetic pass rate = 0.014. The contract-specific reference-to-floor separation is 0.058 (95% CI, 0.036–0.085), below the operational minimum of 0.30, so Table S1 reports NR. Three residue-mapping variants produce comparable geometry-failure counts, arguing against adapter-specific trimming as the cause.

### Supplementary Note 3. BioKinema, the second majority-Layer-4-valid ordered generator

BioKinema[20] is an all-atom diffusion generator designed to reduce autoregressive geometric blow-up. It enters the scorecard in its intended multiple-sequence-alignment configuration and is the second ordered generator after ConfRover to reach majority Layer-4 validity. Its 10-ns temporal contract is range-limited, so Table S1 reports NR.

**Intended configuration.** Every one of the 81 submissions uses a non-single-sequence alignment. Alignment depth ranges from 8 to 12,770 sequences (median 1,315), confirming that the intended MSA-conditioned path was active throughout the evaluation.

**Scores.** BioKinema covers 81 of the 82 proteins and reaches Layer-2 ensemble fidelity 0.639; all 81 trajectories pass Layer 4. The raw 10-ns diagnostics are temporal Layer 3 = 0.129 (95% CI, 0.083–0.177) and post-gate kinetic pass rate = 0.062 (0.012–0.123). The contract-specific reference-to-floor separation is 0.053 (0.032–0.076), so the post-gate kinetic pass is NR. BioKinema therefore supplies a second majority-valid ordered architecture, but not a second resolved kinetic row.

**Matched-subset comparison with ConfRover.** On 72 shared proteins, raw post-gate kinetic pass rates are 0.069 for BioKinema and 0.250 for ConfRover, with Layer-4 validity 1.000 and 0.778. Because BioKinema is range-limited, the interpretable comparison is geometric: BioKinema is more often Layer-4 valid.

**Training-corpus overlap.** BioKinema's checkpoint was trained on a corpus that includes mdCATH. Screening the 82 evaluation chains against 5,286 mdCATH domain chains flags 29 at 30% identity and 50% query coverage, including six at 90% identity or higher. Restricting the analysis to the 52 covered unflagged chains leaves Layer-4 validity at 1.000. The raw post-gate diagnostic is 0.038, while the 10-ns contract remains range-limited.

**Robustness controls.** The threshold sweep, leave-one-protein-out jackknife, common-coverage analysis and sequence-overlap sensitivity retain BioKinema's raw values for diagnostic completeness; none upgrades the contract to a reportable kinetic-pass estimate. BioKinema is a preprint model.

### Supplementary Note 4. The four layers and the metric definitions

Every submission is scored on four layers against the reference MD; the definitions below are those implemented in the released software, and no metric is computed differently for any model.

**Layer 1 (frame realism).** A single-frame sanity check based on virtual Cα–Cα bond geometry and local geometry. On equilibrium MD frames every well-formed frame passes, so Layer 1 is saturated by construction (100% SATURATED across the scorecard) and is not expected to rank plausible generators; it remains explicit so malformed frames fail before ensemble or temporal interpretation.

**Layer 2 (ensemble).** Order-invariant agreement with the equilibrium ensemble: free-energy-surface and contact-distribution divergences and a per-mode displacement 2-Wasserstein distance. These are the quantities an equilibrium generator is meant to reproduce, and are the only layer on which an unordered ensemble submission is scored positively.

**Layer 3 (kinetics), evaluated only for `ordered` submissions.** Eight class-B metrics: time-lagged autocorrelation (`tacf_w1`), VAMP-2 (`vamp2_reldiff`), implied-timescale correlation and Wasserstein (`timescale_logcorr`, `timescale_logw1`), directional predictability (`dir_predict_r2_absdiff`), transition-matrix L1 (`transition_matrix_l1`), mean-first-passage log-Wasserstein (`mfpt_logw1`), and one order-invariant per-mode displacement Wasserstein (`anm_disp_w1`). The first seven are the `TEMPORAL_METRICS` on which the post-gate kinetic endpoint votes; the eighth is informative about a submission but order-invariant, so it is excluded from that vote (an unordered ensemble can legitimately reproduce it). A reader recomputing the Layer-3 *layer score* aggregates all eight; the legacy `realdyn` field uses only the seven time-ordered metrics before applying Layer 4.

**Layer 4 (geometric and fluctuation admissibility).** Five guardrails: steric-clash pair rate, virtual Cα–Cα bad-bond fraction, Ramachandran outlier fraction (only when backbone atoms are supplied), a late-versus-early drift ratio (ordered only) and RMSF spread. The steric rate is the number of eligible non-adjacent Cα pair-frame observations below 4.0 Å divided by the total number of eligible pair-frame observations. Layer 4 has **no class-specific operational reference or single scalar ceiling**; each guardrail defines its own pass region. The clash, bond, Ramachandran and drift guardrails each fire only when a submission exceeds the *larger* of a fixed absolute ceiling and its own protein's reference-replicate envelope inflated by a margin, so a submission is called invalid only when it is bad by both standards at once. The implemented constants are:

| guardrail | absolute ceiling | reference-envelope margin |
|---|---|---|
| steric clash pair rate | 0.000758 | ref-max × (1 + 0.5) |
| bad Cα–Cα bond fraction | 0.02 | ref-max × (1 + 0.5) |
| drift ratio (late/early) | 2.0 | ref-max × 1.5 |
| Ramachandran outlier fraction | NA | ref-max + 0.05 |
| RMSF spread (two-sided) | < 0.5 × ref-min (TOO_COLD); > 2.0 × ref-max (TOO_HOT) | NA |

The clash test excludes sequence separations ≤ 2 and flags Cα–Cα distances below 4.0 Å. Its pair-normalised denominator prevents protein length from inflating the statistic merely by increasing the number of possible contacts. The physical virtual-bond range is 3.0–4.5 Å. Status is assigned by a fixed precedence: local geometry (clash, bond, Ramachandran) is tested first and overrides everything, so a trajectory that fails it is INVALID_GEOMETRY regardless of what else is wrong; only a locally intact trajectory can be TOO_HOT_OR_DIVERGED, then TOO_COLD, else VALID. Radius of gyration is **not** a Layer-4 metric; it is a symmetric quality-control diagnostic for collapse and blow-up, and neither flag automatically excludes a contract-valid submission.

---

### Supplementary Note 5. Anchor construction and the per-class ruler

For each metric on each protein the protocol computes four error values (lower is better) and reads the model's position between them:

- **T, trivial floor.** A surrogate with the target property removed: an i.i.d. resample of the reference frames for kinetics (correct marginal, no temporal order), a static reference structure for the equilibrium layers.
- **O, class-specific operational reference.** A comparator chosen **per metric class**, not a theoretical maximum: an i.i.d. equilibrium resample for the Layer-1–2 equilibrium metrics; a per-mode Ornstein–Uhlenbeck comparator fitted on the reference for most Layer-3 kinetic metrics; a held-out real MD replicate for the transition and mean-first-passage metrics, which a single-basin OU process structurally cannot represent. A richer model may outperform O.
- **S, replicate noise floor.** The agreement one real replicate achieves with another, pooled over the fit replicates as a worst-case bound (the maximum for ATLAS K ≤ 3 replicates, the 90th percentile once K ≥ 4 as on mdCATH, so a single outlier replicate cannot drive a protein to SATURATED).
- **M, model.** The submission under test.

A single universal ruler would systematically mislabel one metric class or the other: using the kinetic OU oracle on the equilibrium layers marks flexible, anharmonic proteins DATA-LIMITED because a Gaussian-around-the-mean surrogate cannot span their ensemble, and using an i.i.d. equilibrium resample on the transition metrics understates what a real replicate can reach. The verdict ladder is then, with `sat_tol = 0.1`, `learn_tol = 0.15` and `triv_gap = T − S`:

1. `triv_gap ≤ 0.1 × scale` → **SATURATED** (the trivial surrogate already reaches the noise floor; the metric cannot discriminate)
2. else `ou_closure = (T − O)/triv_gap < 0.15` → **DATA-LIMITED** (the class-specific reference does not close the gap from trivial under this sampling)
3. else `model_closure = (T − M)/triv_gap ≥ 0.6 × ou_closure` → **MODEL-OK**
4. else `model_closure < 0.15` → **MODEL-FAILS**
5. else → **MODEL-LIMITED**

The production order is part of the frozen implementation. A deterministic sensitivity replay tested MODEL-FAILS before MODEL-OK for every stored metric cell while keeping all closures and thresholds fixed. It changed 0 cells and no kinetic-pass row, so the potentially overlapping weak-reference region is unoccupied on this board. Distinct from all four metric values, a held-out MD replicate is also scored as an ordinary submission: that row is the oracle a reader compares models against and the source of the headline 0.95, and it is not the same object as the per-metric operational reference O.

---

### Supplementary Note 6. Robustness of the headline

Robustness audits bound the ways the trivial-to-oracle bracket or an intermediate model value could depend on analyst choice; three are plotted in Extended Data Fig. 4 and the complete results accompany the public release.

**Pass-fraction-threshold sweep.** The post-gate kinetic endpoint thresholds a protein's informative Layer-3 pass fraction at 0.5. Sweeping it over {0.3, 0.4, 0.5, 0.6, 0.7} preserves the oracle-to-floor bracket but not the ordering of intermediate values. The BioKinema, ProTDyn and MarS-FM columns below are raw diagnostics from unresolved contracts, not reportable post-gate kinetic estimates:

| threshold | oracle | tica_ou | msm | ConfRover | MDGen | BioKinema | MarS-FM | ProTDyn | i.i.d. |
|---|---|---|---|---|---|---|---|---|---|
| 0.3 | 0.988 | 0.378 | 0.439 | 0.397 | 0.134 | 0.235 | 0.000 | 0.027 | 0.000 |
| 0.4 | 0.963 | 0.378 | 0.378 | 0.315 | 0.134 | 0.086 | 0.000 | 0.014 | 0.000 |
| 0.5 | 0.951 | 0.378 | 0.317 | 0.247 | 0.134 | 0.062 | 0.000 | 0.014 | 0.000 |
| 0.6 | 0.866 | 0.317 | 0.159 | 0.110 | 0.098 | 0.012 | 0.000 | 0.014 | 0.000 |
| 0.7 | 0.793 | 0.268 | 0.085 | 0.027 | 0.061 | 0.000 | 0.000 | 0.014 | 0.000 |

The oracle stays at least 0.79 and the i.i.d. floor remains 0.00. BioKinema reaches the floor at 0.7, and ConfRover and MDGen exchange order; no threshold-invariant deep-model ranking is claimed.

**Contract-specific temporal interpretability.** A fail-closed matrix generated one oracle and 20 independent time-destroyed floors for each ordered contract (105 cells total). Every report passed protein-level checks of aligned frame count, effective interval and replicate roles before analysis. The post hoc reporting gate requires an oracle-minus-floor separation of at least 0.30 with a 95% interval excluding zero and median support of at least three informative temporal metrics. Results are:

| model contract | interval | reference frames | median informative metrics | oracle | floor mean | oracle − floor (95% CI) | status |
|---|---|---|---|---|---|---|---|
| ConfRover | 1 ns | 101 | 4 | 0.959 | 0.003 | 0.956 (0.903–0.996) | resolved |
| MDGen | 1 ns | 101 | 4 | 0.927 | 0.005 | 0.921 (0.859–0.971) | resolved |
| BioKinema | 10 ns | 11 | 3 | 1.000 | 0.947 | 0.053 (0.032–0.076) | range-limited |
| ProTDyn | 10 ns | 11 | 3 | 1.000 | 0.942 | 0.058 (0.036–0.085) | range-limited |
| MarS-FM | ~50 ns | 3 | 1 | 0.988 | 0.954 | 0.034 (0.018–0.056) | support-limited |

Excluding MSM metrics when their effective state count is below five gives the same status for every contract. The three coarse-contract gaps are non-zero but operationally insufficient for stable normalization. They therefore receive NR in the formal kinetic-pass field; their raw post-gate values remain below for provenance. BioKinema and ProTDyn share the same 10-ns reference construction and are not independent demonstrations of a universal resolution effect.

**Contract-reportability grid.** Because the 0.30 range and median-three-metric requirements are post hoc operational criteria, we varied the minimum oracle-to-floor gap over 0.20, 0.25, 0.30, 0.35 and 0.40 and the minimum median informative-metric count over 2, 3 and 4, always requiring the gap interval to exclude zero. ConfRover and MDGen are resolved in all 15 combinations. BioKinema, ProTDyn and MarS-FM are unresolved in all 15; only the descriptive label between range-limited and support-limited changes when four metrics are required. This grid tests reportability labels and does not recompute trajectories or verdicts.

**Per-protein support sensitivity.** Contract reportability uses the median number of informative temporal metrics, whereas the post-gate denominator includes any protein with at least one. We therefore recomputed the four resolved rows after requiring at least one, two, three or four informative temporal metrics per protein. Proteins below the stated minimum were excluded before 2,000 protein-bootstrap draws (seed 0). No row contained a protein with only one informative metric. Requiring three removed two proteins from the replicate and i.i.d. rows, three from ConfRover and two from MDGen, with little change in the rates:

| minimum metrics per protein | replicate reference | i.i.d. floor | ConfRover | MDGen |
|---|---|---|---|---|
| 1 | 0.951 (0.902–0.988; n=82) | 0.000 (0.000–0.000; n=82) | 0.247 (0.151–0.342; n=73) | 0.134 (0.061–0.220; n=82) |

| minimum metrics per protein | replicate reference | i.i.d. floor | ConfRover | MDGen |
|---|---|---|---|---|
| 2 | 0.951 (0.902–0.988; n=82) | 0.000 (0.000–0.000; n=82) | 0.247 (0.151–0.356; n=73) | 0.134 (0.061–0.220; n=82) |
| 3 | 0.950 (0.900–0.988; n=80) | 0.000 (0.000–0.000; n=80) | 0.257 (0.157–0.357; n=70) | 0.138 (0.062–0.225; n=80) |
| 4 | 0.970 (0.925–1.000; n=67) | 0.000 (0.000–0.000; n=67) | 0.259 (0.148–0.370; n=54) | 0.091 (0.030–0.167; n=66) |

The support distributions and plotted values accompany the public release. The minimum-three analysis addresses the contract's own support criterion; the minimum-four row is a stricter diagnostic that removes 16–19 proteins from the model rows.

**Replicate-referenced operational ruler.** The production protocol uses metric-specific references, chiefly a fitted Ornstein-Uhlenbeck process for temporal relaxation metrics and a held-out replicate for transition metrics. In a post hoc sensitivity analysis, all seven genuinely temporal metrics instead used the held-out replicate as their operational reference. The time-destroyed floor, temporal raw quantities, non-temporal metrics and Layer 4 were held fixed. The replicate and i.i.d. controls remained at post-gate kinetic pass rates of 1.000 and 0.000. ConfRover changed from 0.247 to 0.397 (paired difference +0.151; 95% CI, +0.055 to +0.247), while MDGen changed from 0.134 to 0.122 (-0.012; -0.037 to 0.000). The audit is floor-coupled rather than fully fit-decoupled and does not replace the frozen board. It shows that the anchor bracket is stable but an intermediate rate depends on the operational reference.

**Leave-one-protein-out jackknife.** Re-scoring the post-gate calculation with each protein held out moves no row by more than 0.014 (oracle 0.012, ConfRover 0.014 and MDGen 0.012; the unresolved ProTDyn and BioKinema raw diagnostics move by 0.014 and 0.013; the i.i.d. floor and MarS-FM remain 0.000), so the anchor bracket is not driven by one protein.

**Leave-one-replicate-out jackknife.** Re-scoring every protein with each of its three MD replicates held out as the evaluation target gives held-out MD post-gate kinetic pass rates of 0.976, 0.976 and 0.951 (mean 0.968, swing 0.024), while the i.i.d. equilibrium control is 0.00 at every hold-out and the reference replicate is Layer-4 valid on 82/82 in all three configurations. The headline 0.95 is the lowest of the three, i.e. the conservative end.

**Decoupled-oracle control.** Removing the shared replicate entirely, with three distinct replicates used for informative-set selection, oracle prediction and evaluation, returns an oracle kinetic-pass rate of 0.963, marginally above the coupled 0.951 (Δ +0.012; the i.i.d. equilibrium control remains 0.00). The informative-set selection therefore does not inflate the ceiling; if anything the standard coupled scoring is slightly conservative.

**Layer-4 guardrail sweep.** The Layer-4 status of a protein is fixed by four tunable constants: the envelope tolerance applied to the clash and virtual-bond statistics, the multiplier on the reference-calibrated absolute clash ceiling, the drift envelope, and the RMSF too-cold and too-hot fractions. Because the per-protein reports store every raw guardrail quantity alongside its reference envelope, the status can be replayed exactly without re-scoring any trajectory. At the frozen defaults the replay reproduces all twelve Layer-4 rows of the board to three decimals, which is the precondition for reading the sweep below.

| row | ceiling 0.25× | ceiling 1× | ceiling 4× | cold 0.3 | cold 0.5 | cold 0.7 |
|---|---|---|---|---|---|---|
| classical_md | 1.000 | 1.000 | 1.000 | 1.000 | 1.000 | 1.000 |
| tica_ou | 0.378 | 0.378 | 0.378 | 0.378 | 0.378 | 0.341 |
| msm_generative | 1.000 | 1.000 | 1.000 | 1.000 | 1.000 | 0.927 |
| iid_equilibrium | 1.000 | 1.000 | 1.000 | 1.000 | 1.000 | 1.000 |
| BioEmu | 0.585 | 0.707 | 0.720 | 0.720 | 0.707 | 0.707 |
| P2DFlow | 0.923 | 0.923 | 0.923 | 0.923 | 0.923 | 0.892 |
| Str2Str | 0.134 | 0.159 | 0.159 | 0.159 | 0.159 | 0.159 |
| ConfRover | 0.658 | 0.781 | 0.836 | 0.849 | 0.781 | 0.452 |
| MDGen | 0.000 | 0.146 | 0.171 | 0.146 | 0.146 | 0.146 |
| MarS-FM | 0.000 | 0.000 | 0.000 | 0.000 | 0.000 | 0.000 |
| ProTDyn | 0.014 | 0.041 | 0.041 | 0.041 | 0.041 | 0.041 |
| BioKinema | 1.000 | 1.000 | 1.000 | 1.000 | 1.000 | 0.926 |

Columns vary one constant with the others held at their frozen values; the 1× and 0.5 columns are both the frozen setting. The envelope tolerance is nearly inert over 0.25–1.5 (only BioEmu moves, 0.707 to 0.720 at 1.5) and the drift envelope moves no row at all, so those two constants are omitted from the table and reported in the JSON. Rescaling the clash ceiling over a factor of sixteen moves absolute values but leaves the five ordered rows in the same order, BioKinema above ConfRover above MDGen above ProTDyn above MarS-FM. The too-cold fraction is the one constant that changes a stated conclusion: ConfRover retains majority Layer-4 validity at 0.6 (0.630) but not at 0.7 (0.452), so its majority status is a claim about generated fluctuation amplitude at the frozen tolerance, not a threshold-free property. BioKinema does not depend on that choice, remaining at or above 0.926 across the whole range. The too-hot fraction acts only on the ensemble rows and on MDGen (0.098 at 1.5 to 0.305 at 3.0) and leaves the anchors and the ordered ranking unchanged.

---

**Sequence-overlap audit (post hoc).** The frozen protocol shipped a k-mer screen; because k-mer Jaccard is a coarse proxy for sequence identity, we additionally ran an upgraded post hoc alignment-based audit with MMseqs2 at sensitivity 7.5, searching the 82 evaluation chains against the 1,266 ATLAS training chains. A protein is flagged at ≥30% identity and ≥50% query coverage, coverage being aligned query residues over the evaluation sequence length. The coverage requirement was added after inspection showed identity alone admits short local matches, so it is a post hoc filter rather than a pre-registered threshold, and the audit is separate from the scoring protocol, which was frozen before promotion and is unchanged by it. The audit flags 10 of 82 proteins, a strict superset of the 4 the k-mer screen flags, leaving an alignment-screened subset of 72 (Supplementary Table S8 gives the threshold grid).

Because rows differ in coverage, the sensitivity analysis is recomputed on the 72-protein intersection shared by the five ordered deep models and four kinetic anchors, of which 62 are unflagged; a protein-level paired bootstrap (2,000 resamples) gives the differences below. Every interval includes zero. Values for unresolved contracts are retained as raw sensitivity diagnostics and are not upgraded to reportable post-gate kinetic estimates.

| row | common (n = 72) | screened common (n = 62) | paired Δ | 95% CI |
|---|---|---|---|---|
| oracle | 0.972 | 0.968 | −0.004 | −0.013 to 0.000 |
| MSM | 0.361 | 0.371 | +0.010 | −0.034 to +0.053 |
| OU | 0.319 | 0.290 | −0.029 | −0.081 to +0.017 |
| ConfRover | 0.250 | 0.258 | +0.008 | −0.033 to +0.044 |
| MDGen | 0.083 | 0.048 | −0.036 | −0.084 to +0.004 |
| BioKinema | 0.069 | 0.048 | −0.021 | −0.065 to +0.009 |
| MarS-FM | 0.000 | 0.000 | 0.000 | 0.000 to 0.000 |
| ProTDyn | 0.014 | 0.016 | +0.002 | 0.000 to +0.009 |
| i.i.d. floor | 0.000 | 0.000 | 0.000 | 0.000 to 0.000 |

The oracle-to-floor separation changes by −0.004 (95% CI, −0.013 to 0.000). Being unflagged means only that no overlap above the operational threshold was detected against this auditable corpus; it is not evidence of the absence of homology or of training-set overlap.

### Supplementary Note 7. Verdict census and metric saturation

Pooled over all thirteen rows, 13,791 metric-by-protein cells receive a verdict and 528 are not applicable by task, giving a 14,319-cell grid.

**DATA-LIMITED = 1.9% (267 of 13,791 scored cells).** Among non-saturated cells the share is 3.0% (267 of 8,998). Unresolved reference signal is uncommon among discriminating cells; this does not prove that no richer model could learn such a cell.

**SATURATED = 29.5% (3,762 of 12,760 scored cells outside the Layer-1 sanity check).** On these cells the trivial surrogate already reaches the replicate noise floor. The share varies among native-task rows because their time intervals leave different temporal metrics informative, so it is reported rather than treated as model-invariant.

---

### Supplementary Note 8. Pre-specified ConfRover failure stratification

ConfRover, the QC-passing ordered model available when the analysis plan was fixed, was pre-specified for stratification of its failures against three protein descriptors: chain length, backbone flexibility (mean Cα RMSF) and collective motion (ANM $r^2$).

**No descriptor stratifies ConfRover's failures.** None is significantly associated with the temporal Layer-3 pass rate, Layer-4 admissibility or post-gate kinetic pass (Spearman $|\rho| \le 0.20$, all $P \ge 0.086$; n = 73, two-sided): Layer-4 admissibility versus length is $\rho = +0.14$ ($P = 0.23$), versus collectivity $\rho = +0.16$ ($P = 0.18$) and versus flexibility $\rho = -0.11$ ($P = 0.34$); post-gate kinetic pass versus length is $\rho = +0.16$ ($P = 0.16$). The failures are therefore not explained by a simple structural descriptor and cannot be dismissed as an effect of protein size or flexibility alone.

**Reference resolvability holds in every stratum.** The replicate oracle's kinetic-pass rate stays at 0.89–1.00 across every tertile of every descriptor, so no stratum is left unresolved by the reference data; ConfRover's failures are the model's, not absent reference signal. Flexibility and collectivity are strongly anticorrelated ($\rho = -0.66$), whereas length shows no detected monotonic association with either ($|\rho| \le 0.06$); this does not establish independence. This is a within-model diagnostic for ConfRover, not a claim about ordered models in general.

---

### Supplementary Note 9. Synthetic ordering of learned ensembles

The frame-shuffle control in the main analysis uses authentic MD and therefore isolates temporal order exactly. We additionally asked whether the same measurement failure appears in outputs learned by unordered conformational generators. For BioEmu, P2DFlow and Str2Str, a fixed set of 50 generated frames per protein was assigned ten independently seeded random orders. One BioEmu protein with only ten available frames was excluded. The scored coverage was therefore 81, 65 and 82 proteins. Direct coordinate comparison verified that every seed used the same selected frames and changed only their order; no selected model frame matched the held-out reference coordinate fingerprint.

These are synthetic-order stress tests, not kinetic evaluations. The source models remain unordered by task, no synthetic interval is interpreted as physical time, and their scorecard kinetic entries remain NA. The 50-frame model support is also shorter than the standard 101-frame reference support, so the temporal values are reported only as within-model diagnostics and are not normalised against the reference.

| model | proteins | L2 ensemble, all orders | temporal L3, mean | temporal L3, range over 10 orders | direct reference-coordinate matches |
|---|---|---|---|---|---|
| BioEmu | 81 | 0.455 | 0.096 | 0.065–0.111 | 0 |
| P2DFlow | 65 | 0.672 | 0.130 | 0.099–0.158 | 0 |
| Str2Str | 82 | 0.227 | 0.103 | 0.087–0.116 | 0 |

Layer-2 fidelity is exactly invariant across synthetic orders, while only a small fraction of order-sensitive tests pass. This extends the controlled MD permutation result to learned conformational outputs without reclassifying ensemble generators as trajectory models or assigning them kinetic credit.

---

### Supplementary Note 10. Reference-only temporal-resolution and support audit

The native-contract audit showed that 1-ns ConfRover and MDGen outputs are reportable whereas the tested 10-ns and approximately 50-ns contracts are not. To separate temporal interval from observation horizon, we repeated the leak-free A/B/C reference-support construction on all 82 ATLAS proteins without scoring a learned model. Replica C remained the evaluation target, replicas A and B formed the fit and i.i.d.-floor pool, and B served as the reference stand-in. Every condition used 20 independent floor seeds and 2,000 hierarchical bootstrap draws.

At a fixed 100-ns horizon, the oracle-minus-floor gap decreased from 0.961 at 1 ns to 0.767 at 2 ns, 0.535 at 4 ns, 0.472 at 5 ns and 0.023–0.063 at 10–50 ns. The 4- and 5-ns conditions exceeded the numerical gap criterion but retained a median of only two informative temporal metrics; only 1 and 2 ns therefore met both reportability criteria. In a complementary fixed-support arm, 11 frames were retained at intervals of 1, 2, 4, 5 and 10 ns. Every condition was unresolved, with gaps of 0.039–0.061 despite a median of three informative metrics. The result shows that the tested measurement requires both adequate temporal resolution and adequate observation length. The grid is post hoc and reference-only, so it neither ranks learned models nor establishes a universal cutoff for other datasets or observables.

---

### Supplementary Note 11. Prospective Lockbox-36 validation

Lockbox-36 tests the frozen measurement logic on proteins selected after the evaluated model checkpoints were fixed. The Broad arm contains 36 soluble monomeric proteins of 60–300 residues, each with three independent 200-ns CHARMM36m simulations. Eight proteins were selected for the Deep arm before any Broad kinetic result was inspected, and all three replicas were extended to 1 μs. Candidate identities, replicate roles, model contracts, endpoints, exclusions, bootstrap rules and gate thresholds were fixed before scoring. ConfRover and MDGen are the primary 1-ns ordered contracts, BioEmu is an unordered applicability control and BioKinema is a secondary 10-ns contract audit.

An initial extraction was invalidated because wrapped periodic coordinates broke local Cα chains and inflated reference fluctuation envelopes. No score from that extraction is used. Whole molecules were reconstructed across the periodic box, the corrected extraction and analysis code were frozen before any corrected score was inspected, and all 36 Broad and eight Deep reports subsequently passed completeness and point-estimate verification.

| prospective arm | reference | i.i.d. floor | ConfRover | MDGen | paired ConfRover minus MDGen |
|---|---|---|---|---|---|
| Broad, 36 proteins, 200 ns | 1.000 | 0.000 | 0.111 (0.028–0.222) | 0.583 (0.417–0.750) | −0.472 (−0.667 to −0.250) |
| Broad, 27 unflagged proteins | 1.000 | 0.000 | 0.148 (0.037–0.296) | 0.593 (0.407–0.778) | −0.444 (−0.667 to −0.185) |
| Deep, 8 proteins, 1 μs | 0.963 | 0.000 | 0.075 (0.000–0.200) | 0.462 (0.188–0.700) | −0.388 (−0.650 to −0.150) |

Broad values are post-gate kinetic-pass fractions. Deep values average ten non-overlapping 100-frame windows within each protein; the paired column is the window-averaged pass difference. Normalising the Deep model rates by the reference-to-floor range gives anchor-gap attainments of 0.078 (0.000–0.213) for ConfRover and 0.481 (0.192–0.727) for MDGen, with no non-positive denominator among 2,000 bootstrap draws. In Broad, MDGen exceeds ConfRover in both temporal-majority passes (0.778 versus 0.306) and Layer-4 validity (0.694 versus 0.361). The prospective reversal therefore reflects both temporal and geometric evidence, not a gate-only effect. BioEmu reaches Layer-2 fidelity 0.813 and Layer-4 validity 0.667 on all 36 proteins; its temporal endpoint remains NA by task.

The full frozen-input sequence screen searched all 36 proteins against the disclosed ATLAS and mdCATH corpora at at least 30% identity and at least 50% query coverage. Nine proteins were flagged. Their removal leaves the model direction unchanged in Broad and in the six-protein unflagged Deep subset. This sensitivity addresses the disclosed corpora and does not establish absence of overlap with unavailable training data.

BioKinema has 32 of 36 technically evaluable Broad outputs and seven of eight Deep outputs; the remaining outputs are technical NA rather than zero scores. At the 10-ns interval, the 200-ns reference supplies 21 frames, median temporal support 2.0 and only 5 of 15 resolved reportability-grid settings. The 1-μs Deep reference supplies 101 frames, median support 5.5 and 15 of 15 resolved settings. The primary generated sample passes on one of seven evaluable Deep proteins, whereas each of four additional samples passes on none (Fig. 4c). Longer observation therefore makes the contract measurable without converting greater temporal support into model success. The complete source data and audit records accompany the public evidence package.

**Metric-family sensitivity (post hoc).** The frozen Lockbox protocol named sensitivity across metric families as a secondary analysis, but did not freeze the family partition. We therefore formalised four complete groups after scoring and did not search metric combinations: timescale (correlation and Wasserstein), autocorrelation, transition matrix plus MFPT, and directional prediction plus VAMP. Each row below removes one group, recomputes temporal majority and reportability from stored verdicts, retains native-rollout Layer 4 and applies 2,000 paired protein-bootstrap draws with crossed resampling of 20 floor seeds.

| temporal panel | Broad reference−floor | Broad support | Broad ConfRover−MDGen (95% CI) | Deep reference−floor | Deep support | Deep ConfRover−MDGen (95% CI) |
|---|---|---|---|---|---|---|
| all seven metrics | 1.000 | 6.00 | −0.472 (−0.667 to −0.278) | 0.962 | 5.00 | −0.387 (−0.650 to −0.125) |
| without timescale | 0.972 | 4.00 | −0.417 (−0.611 to −0.222) | 0.962 | 4.00 | −0.287 (−0.512 to −0.075) |
| without autocorrelation | 0.994 | 5.00 | −0.500 (−0.694 to −0.306) | 0.950 | 4.75 | −0.425 (−0.725 to −0.100) |
| without transition/MFPT | 0.950 | 4.00 | −0.500 (−0.667 to −0.306) | 0.938 | 3.25 | −0.500 (−0.837 to −0.150) |
| without directional/VAMP | 0.992 | 4.00 | −0.306 (−0.500 to −0.111) | 0.962 | 3.75 | −0.275 (−0.500 to −0.050) |

Both reference contracts remain resolved after every family removal, and every paired model interval remains below zero. The result shows that neither the prospective anchor separation nor the model-order reversal is carried by one temporal family; it does not define a minimal metric panel. The measured ATLAS and Lockbox support conditions are additionally assembled without interpolation; these discrete points do not define a continuous phase boundary.

### Supplementary Note 12. Stationary-matched transition-rate perturbation control

Frame shuffling proves that ensemble agreement can survive complete loss of order. We next constructed a harder adversarial control in which every arm contains the exact same authentic coordinates in the same multiplicities but traverses their microstates under a different transition operator. For each of the 82 ATLAS proteins, a reversible microstate Markov model was fitted to the two fit replicates. The held-out third replicate was excluded from state-space fitting, transition estimation and frame emissions. Five operator arms were labelled by nominal dynamical-speed factors of 0.25, 0.5, 1, 2 and 4 while preserving the fitted stationary distribution.

Within each protein and seed, stationary state counts were rounded once and one multiset of authentic fit-replicate frames was drawn. A conditioned path sampler then reordered that exact coordinate multiset under each transition operator. All 410 protein-seed groups passed byte-level coordinate-multiset equality across rates, all 2,050 requested cells completed and the maximum stationary residual was $5.56 \times 10^{-17}$. Five fixed seeds were averaged within protein before 2,000 protein-bootstrap resamples.

| nominal speed factor | adjacent state changes | Layer 2 | temporal Layer 3 (95% CI) | Layer 4 | post-gate kinetic pass (95% CI) |
|---|---|---|---|---|---|
| 0.25 | 0.252 | 0.990 | 0.648 (0.607–0.686) | 0.998 | 0.815 (0.744–0.876) |
| 0.50 | 0.319 | 0.990 | 0.621 (0.579–0.661) | 1.000 | 0.776 (0.700–0.841) |
| 1.00 | 0.435 | 0.990 | 0.581 (0.539–0.622) | 0.998 | 0.729 (0.649–0.805) |
| 2.00 | 0.529 | 0.990 | 0.519 (0.482–0.556) | 1.000 | 0.644 (0.561–0.724) |
| 4.00 | 0.615 | 0.990 | 0.445 (0.407–0.481) | 0.998 | 0.546 (0.466–0.627) |

Systematic acceleration reduced both temporal readouts while leaving the coordinate-defined measurements fixed. Relative to the 1-fold arm, the temporal Layer-3 changes were −0.062 (95% CI, −0.080 to −0.044) at 2-fold and −0.136 (−0.164 to −0.107) at 4-fold. The corresponding post-gate changes were −0.085 (−0.124 to −0.049) and −0.183 (−0.239 to −0.124). Layer 2 was exactly invariant and the largest Layer-4 point change was 0.0024.

The slower arms increased rather than decreased the temporal scores: 0.25-fold versus 1-fold changed Layer 3 by +0.067 (0.046–0.090) and post-gate kinetic pass by +0.085 (0.046–0.129). The fitted 1-fold Markov chain is not a physical-rate oracle, so this asymmetry does not identify an optimal rate. It shows that Dynbench detects a controlled systematic acceleration without relying on altered state populations, novel coordinates or geometric failure. The control is an audit and does not replace a learned-model scorecard row.

### Supplementary Table S1. Contract-audited scorecard (ATLAS-82; 95% protein-cluster bootstrap CIs)

Layer 3 is the temporal-only raw statistic used by the post-gate calculation and is not applicable to unordered ensemble submissions. The final column is the post-gate kinetic pass rate and is reported only when the contract-specific oracle-to-floor audit is resolved. AlphaFlow's Layer-4 value is not reported because its complete coordinate batch was unavailable.

| model | type | contract status | n | L2 | temporal L3 | L4 | post-gate kinetic pass |
|---|---|---|---|---|---|---|---|
| classical_md | oracle (real MD replicate) | 1 ns, resolved | 82 | 0.881 (0.850–0.909) | 0.825 (0.784–0.863) | 1.000 (1.000–1.000) | **0.951 (0.890–0.988)** |
| tica_ou | surrogate (per-mode OU) | 1 ns, resolved | 82 | 0.845 (0.809–0.880) | 0.846 (0.812–0.883) | 0.378 (0.280–0.488) | 0.378 (0.280–0.488) |
| msm_generative | surrogate (MSM on TICA) | 1 ns, resolved | 82 | 0.810 (0.764–0.858) | 0.295 (0.234–0.357) | 1.000 (1.000–1.000) | 0.317 (0.220–0.427) |
| iid_equilibrium | trivial (i.i.d. resample) | 1 ns, resolved | 82 | **0.99 (0.98–1.00)** | 0.00 (0.00–0.00) | 1.00 (1.00–1.00) | **0.00 (0.00–0.00)** |
| ConfRover | ordered deep model | 1 ns, resolved | 73 | 0.493 (0.450–0.538) | 0.301 (0.255–0.353) | 0.781 (0.671–0.877) | **0.247 (0.151–0.356)** |
| MDGen | ordered deep model | 1 ns, resolved | 82 | 0.206 (0.173–0.242) | 0.643 (0.603–0.685) | 0.146 (0.073–0.220) | 0.134 (0.061–0.207) |
| BioKinema | intended-MSA; training overlap (Note 3) | 10 ns, range-limited | 81 | 0.639 (0.597–0.681) | 0.129 (0.083–0.177), raw | 1.000 (1.000–1.000) | **NR**; raw 0.062 (0.012–0.123) |
| ProTDyn | ordered deep model, QC-failing | 10 ns, range-limited | 74 | 0.146 (0.116–0.180) | 0.238 (0.170–0.311), raw | 0.041 (0.000–0.095) | **NR**; raw 0.014 (0.000–0.041) |
| MarS-FM | ordered deep model | ~50 ns, support-limited | 82 | 0.091 (0.069–0.113) | 0.295 (0.222–0.371), raw | 0.000 (0.000–0.000) | **NR**; raw 0.000 (0.000–0.000) |
| AlphaFlow | ensemble deep model | unordered | 82 | 0.635 (0.570–0.690) | NA-by-task | NR | NA-by-task |
| P2DFlow | ensemble deep model | unordered | 65 | 0.680 (0.613–0.746) | NA-by-task | 0.923 (0.846–0.985) | NA-by-task |
| BioEmu | ensemble deep model | unordered | 82 | 0.480 (0.424–0.539) | NA-by-task | 0.707 (0.610–0.817) | NA-by-task |
| Str2Str | ensemble deep model | unordered | 82 | 0.220 (0.192–0.250) | NA-by-task | 0.159 (0.085–0.244) | NA-by-task |

### Supplementary Table S2. Per-model quality-control cards

Coverage, output contract, geometry diagnostics and Layer-4 status distributions. Radius of gyration is a QC diagnostic rather than a verdict input. NR denotes a Layer-4 result that could not be computed.

| model | n | contract | dominant failure | Rg ratio (med) | Cα–Cα ok (med) | L4: VALID / INVALID_GEOM / TOO_HOT / TOO_COLD |
|---|---|---|---|---|---|---|
| AlphaFlow | 82 | unordered ensemble | NA-by-task (kinetics) | 0.998 | 0.998 | NR |
| P2DFlow | 65 | unordered ensemble | NA-by-task (kinetics) | 1.007 | 1.000 | 60 / 2 / 3 / 0 |
| BioEmu | 82 | unordered ensemble | NA-by-task (kinetics) | 0.988 | 0.992 | 58 / 1 / 22 / 1 |
| Str2Str | 82 | unordered ensemble | over-dispersed geometry | 1.016 | 1.000 | 13 / 18 / 51 / 0 |
| ConfRover | 73 | ordered trajectory | native-task kinetics limited | 0.998 | 0.988 | 57 / 9 / 2 / 5 |
| MDGen | 82 | ordered trajectory | invalid geometry | 1.09 | 0.978 | 12 / 45 / 25 / 0 |
| MarS-FM | 82 | ordered trajectory | geometric blow-up | 5.418 | 0.462 | 0 / 82 / 0 / 0 |
| ProTDyn | 74 | ordered trajectory | invalid geometry | 1.078 | 0.511 | 3 / 59 / 12 / 0 |

| model | n | contract | dominant failure | Rg ratio (med) | Cα–Cα ok (med) | L4: VALID / INVALID_GEOM / TOO_HOT / TOO_COLD |
|---|---|---|---|---|---|---|
| BioKinema | 81 | ordered trajectory, intended-MSA | native-task kinetics limited | NA | 1.000 | 81 / 0 / 0 / 0 |

### Supplementary Table S3. Verdict census (13,791 scored metric-by-protein cells)

| verdict | count | % of scored | in denominator? |
|---|---|---|---|
| MODEL-OK | 4,455 | 32.3% | yes (the only pass) |
| SATURATED | 4,793 | 34.8% | no (leaves the denominator) |
| MODEL-FAILS | 3,049 | 22.1% | yes |
| MODEL-LIMITED | 1,227 | 8.9% | yes (counts against the model) |
| DATA-LIMITED | 267 | 1.9% | yes (counts against the model) |
| **total scored** | **13,791** | 100% | NA |

A further 528 metric cells are not applicable by task, so the full grid is 14,319 cells. Excluding the Layer-1 sanity check, SATURATED is 3,762 of 12,760 = 29.5%. See Supplementary Note 7.

### Supplementary Table S4. Small-peptide positive-control ladders

Baseline ladder on the peptide audit sets (no deep model), on heavy-atom pairwise-distance observables. Temporal Layer 3 is the informative-pass fraction over the seven time-ordered metrics used by the kinetic gate; the separate legacy class-B statistic is not shown. Post-gate kinetic pass is the Layer-4-gated binary verdict. Provenance and the temporal-only recomputation accompany the public release.

| dataset | baseline | L2 | temporal L3 (n informative) | post-gate kinetic pass |
|---|---|---|---|---|
| Timewarp AD (2 traj) | classical_md | 1.00 | 0.67 (6) | 1.00 |
| | msm_generative | 1.00 | 0.00 (6) | 0.00 |
| | tica_ou | 0.80 | 0.00 (6) | 0.00 |
| | iid_equilibrium | 1.00 | 0.00 (6) | 0.00 |
| MDshare AD (3 traj) | classical_md | 1.00 | 1.00 (5) | 1.00 |
| | msm_generative | 1.00 | 1.00 (5) | 1.00 |
| | tica_ou | 0.50 | 0.80 (5) | 1.00 |
| | iid_equilibrium | 1.00 | 0.00 (5) | 0.00 |
| WLALL pentapeptide (25 traj) | classical_md | 1.00 | 1.00 (1) | 1.00 |
| | msm_generative | 1.00 | 1.00 (1) | 1.00 |
| | tica_ou | 0.71 | 1.00 (1) | 1.00 |
| | iid_equilibrium | 1.00 | 0.00 (1) | 0.00 |

### Supplementary Table S5. External protein-set validation

Baseline ladder on two off-ATLAS protein datasets, establishing that the trivial-to-oracle dissociation is not an ATLAS artefact. mdCATH is evaluated at 320 K and a 4-ns interval; PTM-85 contains phospho-serine and three replicates per protein.

| dataset | baseline | L2 | temporal L3 | L4 valid | post-gate kinetic pass |
|---|---|---|---|---|---|
| mdCATH-82 | classical_md | 0.95 | 0.89 | 1.00 | 0.98 |
| | tica_ou | 0.65 | 0.75 | 0.00 | 0.00 |
| | msm_generative | 0.96 | 0.34 | 0.99 | 0.28 |

| dataset | baseline | L2 | temporal L3 | L4 valid | post-gate kinetic pass |
|---|---|---|---|---|---|
| | iid_equilibrium | 0.99 | 0.00 | 1.00 | 0.00 |
| PTM-85 | classical_md | 0.93 | 0.85 | 1.00 | 0.98 |
| | tica_ou | 0.81 | 0.85 | 0.52 | 0.52 |
| | msm_generative | 0.75 | 0.23 | 0.99 | 0.18 |
| | iid_equilibrium | 1.00 | 0.00 | 1.00 | 0.00 |

On both corpora the i.i.d. floor reproduces the ensemble (Layer-2 ≈ 0.99–1.00) yet none of the kinetics (Layer-3 0.00), while the replicate oracle passes the majority of kinetic metrics and the Layer-4 geometric guardrails on every protein, reproducing the ATLAS signature.

The completed ConfRover mdCATH rollout additionally provides a bounded model-level external test. Under a replicate-referenced temporal ruler, its temporal pass fraction is 0.228, compared with 0.997 for held-out MD and 0.000 for the i.i.d. floor; the corresponding post-gate rates are 0.073, 1.000 and 0.000. Protein-paired differences place ConfRover above the floor by +0.073 (95% CI, +0.024 to +0.134) and below the replicate by 0.927 (0.866 to 0.976). The model's Layer-4-valid fraction is 0.220. This is one external ConfRover contract, not an independent multi-model board.

---

### Supplementary Table S6. Audited-dataset provenance

Each external and small-peptide set carries a record of provenance, stride, observable and oracle/floor definitions. Public sources were the mdCATH and Microsoft Timewarp releases and the WLALL and MDshare collections; PTM-85 was generated in house. The mdCATH dataset DOI is 10.1038/s41597-024-04140-z.

| dataset | replicates | sampling | observable | oracle |
|---|---|---|---|---|
| mdCATH-82 | multi-temperature, held-out matched replica | 4-ns interval | Cα (T,L,3) | held-out temperature-matched replicate |
| PTM-85 | 3 per protein | native 100-ps interval | Cα (T,L,3) | held-out replicate |
| WLALL pentapeptide | 25 | stride 25 | heavy-atom distances | cross-replicate (25 trajectories) |
| Timewarp AD | 2 (ad1/ad2) | stride 100 | heavy-atom distances | cross-replicate |
| MDshare AD | 3 | stride 50 | heavy-atom distances | cross-replicate (3 trajectories) |

---

### Supplementary Table S7. Eligibility and availability of ordered protein-dynamics generators

The table records protein-scale ordered-output capability and public availability at the evaluation cutoff of 24 August 2026. `Ref.` is the main-manuscript reference number, or an arXiv/venue identifier for works not in that list. Availability determines whether a method can be evaluated independently; it is not a measure of model quality. Methods published after the scoring freeze are included for field coverage but were not added retrospectively.

**Adjacent experiment-anchored benchmark.** Zhang, Miller and Bowman evaluated AlphaFlow, BioEmu and related methods against experimentally measured cryptic-pocket opening populations and mutation effects[15]. This provides an equilibrium-population test along a defined pocket-opening coordinate and also reports structurally implausible outputs. It does not test an ordered protein trajectory or temporal transition rates, so it is cited as adjacent evidence rather than entered in this ordered-generator census, the capability matrix or the frozen board.

**Availability and scope.**

| Method | Ref. | Protein-scale ordered output | Code public | Weights public |
|---|---|---|---|---|
| ConfRover | 10 | Yes | Yes | Yes |
| MDGen | 9 | Yes (OOD-limited) | Yes | Yes |

| Method | Ref. | Protein-scale ordered output | Code public | Weights public |
|---|---|---|---|---|
| MarS-FM | 11 | Yes | Yes | Yes |
| ProTDyn | 12 | Yes | Yes | Yes |
| BioKinema | 20 | Yes | Yes | Partial (CATH and octapeptide checkpoint; corpus includes mdCATH) |
| Beyond Ensembles (GLDP) | 18 | Yes | Yes | Not evaluated |
| STAR-MD | ICLR 2026; arXiv:2602.02128 | Yes | No (coming soon) | No (coming soon) |
| Pretrained Variational Bridge | arXiv:2602.07588 | Yes | No public release located | No public release located |
| ATMOS | arXiv:2603.17633 | Yes | No public release located | No public release located |
| BioDynaSpec | ICML 2026 | Yes | Yes | No public checkpoint located |
| DynaMode | ICML 2026 GenBio; arXiv:2607.04134 | Yes | Yes | No (available soon) |
| DyneTrion | arXiv:2607.15309 | Yes | Yes | Yes (Hugging Face checkpoint) |
| Timewarp | 22 | No (peptides) | Yes | Yes |
| Score Dynamics | 16 | No (small systems) | Not evaluated | Not evaluated |

#### Evaluation eligibility.

| Method | Contract assessment | Disposition |
|---|---|---|
| ConfRover | Compatible | Post-gate kinetic pass 0.247; 57/73 Layer-4 valid |
| MDGen | Compatible | Post-gate kinetic pass 0.134; Layer-4-gated (12/82 valid) |
| MarS-FM | Compatible; support-limited | Post-gate kinetic pass NR; raw diagnostic 0.000; 0/82 Layer-4 valid |
| ProTDyn | Compatible; range-limited | Post-gate kinetic pass NR; raw diagnostic 0.014; 3/74 Layer-4 valid (Note 2) |
| BioKinema | Compatible in intended-MSA mode (81/82) | Post-gate kinetic pass NR; raw diagnostic 0.062; 81/81 Layer-4 valid; training overlap 29/82 (Note 3) |
| Beyond Ensembles (GLDP) | Published after scoring freeze | Literature context only |
| STAR-MD | No public code or weights at cutoff | Not independently evaluable |
| Pretrained Variational Bridge | No public code or weights at cutoff | Not independently evaluable |
| ATMOS | No public code or weights at cutoff | Not independently evaluable |
| BioDynaSpec | No compatible released checkpoint at cutoff | Not independently evaluable |
| DynaMode | No released checkpoint at cutoff | Not independently evaluable |
| DyneTrion | Published after scoring freeze | Literature context only |
| Timewarp | Peptide-scale task | Outside protein-scale scope |
| Score Dynamics | Small-system task | Outside protein-scale scope |

Five eligible models entered the ordered scorecard. ConfRover and MDGen clear the temporal reference-support gate. BioKinema is geometrically admissible but range-limited at its native 10-ns contract; ProTDyn is range-limited and fails Layer 4; MarS-FM is support-limited. BioKinema additionally overlaps its reported training corpus for 29 of 82 ATLAS chains (Supplementary Note 3).

Methods without released code or checkpoints are recorded as not independently evaluable, not as performance failures. TrajCast is a general molecular and materials dynamics generator rather than a protein conformational-dynamics method. Mac-Diff (ref. 8) is an unordered ensemble generator and therefore lies outside this ordered census; by task, it would not receive a kinetic verdict.

### Supplementary Table S8. Sequence-overlap audit threshold grid (MMseqs2)

Proteins flagged out of 82, and the size of the resulting alignment-screened subset, over identity × query-coverage. The main text uses identity ≥ 0.30 with query coverage ≥ 0.50 (bold). Source data are provided with this paper.

| identity \ query coverage | 0.0 | 0.3 | 0.5 | 0.8 |
|---|---|---|---|---|
| 0.20 | 67 / 15 | 31 / 51 | 18 / 64 | 10 / 72 |
| 0.25 | 67 / 15 | 28 / 54 | 15 / 67 | 8 / 74 |
| 0.30 | 59 / 23 | 19 / 63 | **10 / 72** | 7 / 75 |
| 0.40 | 44 / 38 | 11 / 71 | 8 / 74 | 6 / 76 |
| 0.50 | 28 / 54 | 6 / 76 | 5 / 77 | 3 / 79 |

Cells are flagged / screened-subset size. The paired score analysis at the main-text setting is in Supplementary Note 6; the grid itself is an audit of subset composition, not a claim of score invariance at every cutoff.
